# Transcriptional rewiring of NF-κB by HBZ and Fra-2 drives inflammatory chemoresistance in Tax-negative adult T-cell leukemia/lymphoma

**DOI:** 10.64898/2026.09.30.755641

**Authors:** Julie Tram, Laetitia Marty, Célima Mourouvin, Magali Abrantes, Raymond Césaire, Philippe Hélias, Jean-Michel Mesnard, Véronique Baccini, Jean-Marie Peloponese

## Abstract

Adult T-cell leukemia/lymphoma (ATL) is an aggressive HTLV-1–associated malignancy with poor responses to chemotherapy. Although constitutive NF-κB activation is a hallmark of ATL, its persistence in Tax-negative (Type I) ATL—where Tax is irreversibly silenced—remains unresolved. Here, we identify Fra-2 as a coactivator of NF-κB p65 that redirects p65’s transcriptional activity toward inflammatory genes. Mechanistically, Fra-2 interacts with the Rel homology domain of p65 via its bZIP domain, enhancing p65 DNA-binding activity without altering nuclear localization. The HTLV-1 antisense protein HBZ disrupts Fra-2/JunD AP-1 complexes while stabilizing Fra-2/p65, as shown by co-immunoprecipitation, BiFC, and PICO assays. This rewiring selectively enhances NF-κB-dependent transcription of IL-1 β, which is elevated in ATL patient sera and cells independently of proviral load. Chromatin immunoprecipitation revealed Fra-2 recruitment to NF-κB response elements in the IL1β promoter, an effect amplified by HBZ. Pharmacological inhibition of IL-1 signaling with Anakinra or anti-IL-1β antibodies restored doxorubicin sensitivity in primary ATL cells. Together, these findings establish the Fra-2–NF-κB–IL-1β axis as a driver of chemoresistance in Tax-negative ATL and a therapeutically targetable vulnerability.

## Introduction

Despite advances in therapy, adult T-cell leukemia/lymphoma (ATL) remains an aggressive HTLV-1–associated malignancy characterized by constitutive inflammatory signaling and poor responses to chemotherapy ^1–4^. Recent studies have revealed that ATL is far more heterogeneous than previously appreciated with respect to HTLV-1 viral gene expression. In particular, Nosaka and Matsuoka proposed a classification in which ATL can be divided into three molecular subtypes based on the expression pattern of the viral transactivator Tax^5^. Despite these differences, one feature is remarkably consistent across all subtypes: the continuous expression of the viral antisense protein HBZ, highlighting its central role in maintaining the leukemic phenotype. In a small subset of patients (Type III), Tax remains highly expressed, resembling the latency III program observed in Epstein–Barr virus-associated malignancies^5^. By contrast, nearly half of ATL cases (Type II) retain an intact provirus, including a functional 5′ long terminal repeat (LTR) and an intact *tax* coding sequence, indicating that they preserve the capacity to express Tax. However, some suggest that Tax expression in these tumors is tightly controlled and occurs only transiently, most likely through reversible epigenetic mechanisms controlled by HBZ that balance viral persistence with immune evasion. Indeed, HBZ has been shown to further repress Tax-dependent transcription by competing with Tax for the transcriptional coactivators CBP/p300 ^6,7^, thereby reinforcing viral latency. The remaining ATL cases (Type I) have permanently lost Tax expression through irreversible genetic or epigenetic alterations. These include deletion of the 5′ LTR, inactivating mutations within the *tax* gene, insertion/deletion events, or promoter hypermethylation^8–10^.

While constitutive NF-κB activation is well-documented in ATL ^11–13^, the mechanisms sustaining this pathway in Tax-negative Type I ATL—where Tax is irreversibly silenced— remain poorly understood. Furthermore, the role of AP-1 components in modulating NF-κB activity in ATL has not been systematically explored, despite evidence of crosstalk in other cancers ^14–16^. Here, we investigate whether Fra-2, an AP-1 family member, cooperates with NF-κB to drive inflammatory chemoresistance in ATL, and whether HBZ orchestrates this process.

A major obstacle in the treatment of ATL is the development of resistance to cytotoxic agents, which markedly reduces the efficacy of standard regimens and contributes to disease relapse. Increasing evidence indicates that inflammatory signaling pathways play a central role in leukemic cell survival and therapeutic resistance in ATL^1–4^. Among these pathways, nuclear factor κB (NF-κB) signaling is particularly important ^17–19^. NF-κB transcription factors regulate genes involved in cell survival, proliferation, inflammation, and stress responses, and constitutive NF-κB activation is a hallmark of HTLV-1–transformed cells ^12,20,21^. Notably, NF-κB signaling remains active in most ATL cells despite the frequent absence of Tax expression, indicating that additional mechanisms sustain canonical NF-κB activation in established disease ^12,20,21^. Persistent NF-κB activity promotes the expression of anti-apoptotic proteins such as Bcl-2 and Bcl-xL, enhances cell-cycle progression, and induces the production of cytokines and chemokines that create a tumor-supportive microenvironment characterized by chronic inflammation and immune cell recruitment ^12,20,21^. These processes collectively contribute to leukemic cell survival and intrinsic chemoresistance.

Cytokines mediate communication between malignant cells and their microenvironment and play critical roles in tumor progression and therapeutic response ^17,22–26^. Proinflammatory cytokines can promote cell survival, angiogenesis, and immune modulation ^17,22–26^. Consistent with this concept, we observed that sera from type I ATL patients contained elevated levels of inflammatory cytokines, including IL-1α, IL-1β, IL-2, IL-10, IL-12, TNF-α, and GM-CSF, compared with sera from HTLV-1 asymptomatic carriers, suggesting that dysregulated cytokine production is a prominent feature of the ATL inflammatory microenvironment.

While constitutive NF-κB activation is well-documented in ATL, the mechanisms sustaining this pathway in Tax-negative Type I ATL—where Tax is irreversibly silenced—remain poorly understood. Furthermore, the role of AP-1 components in modulating NF-κB activity in ATL has not been systematically explored, despite evidence of crosstalk in other cancers. Here, we investigate whether Fra-2, an AP-1 family member, cooperates with NF-κB to drive inflammatory chemoresistance in ATL, and whether HBZ orchestrates this process. The molecular mechanisms linking inflammatory signaling to transcriptional regulation in ATL remain incompletely defined. Activator protein 1 (AP-1) transcription factors regulate genes involved in proliferation, differentiation, and stress responses ^27–29^. Functional interactions between AP-1 and NF-κB pathways have been described in inflammatory and malignant contexts ^17,30–32^. However, the contribution of specific AP-1 family members to NF-κB– dependent transcription and chemoresistance in ATL remains poorly characterized. In this study, we identify the AP-1 family member Fra-2 as a regulator of canonical NF-κB signaling in ATL type I. We show that, unlike cFos, Fra-2 interacts with the Rel homology domain of NF-κB p65 through its bZIP region and enhances NF-κB–dependent transcription. Fra-2 promotes transactivation of inflammatory cytokine promoters, including IL-1α, IL-1β, IL-2, TNF-α, and GM-CSF, thereby fostering a microenvironment that supports leukemic cell survival. Chromatin immunoprecipitation analysis demonstrated that Fra-2/p65 complexes preferentially bind NF-κB elements rather than AP-1 sites within these promoters, indicating a functional redirection of Fra-2 toward NF-κB–dependent transcription.

We further demonstrate that the HTLV-1–encoded protein HBZ enhances Fra-2–mediated activation of NF-κB and cytokine gene promoters, suggesting that viral and host transcriptional regulators cooperate to amplify inflammatory signaling in ATL. These findings identify a mechanism by which inflammatory transcriptional programs are sustained in leukemic cells and provide insight into how cytokine-driven signaling contributes to chemoresistance. Understanding these pathways could guide the creation of therapies that block inflammatory survival signals, potentially enhancing treatment outcomes in ATL.

## Materials and Methods

### HTLV-1 Patients

Serum and blood samples were obtained from the Centre de Ressources Biologiques of Martinique (CeRBiM) (BRIF BB-0033-00099) at the Teaching Hospital of Fort-de-France, Martinique, from collections NCT01754311 and NCT01941680. The diagnosis of ATL was based on clinical features, hematological findings, detection of the HTLV-1 provirus in leukemic cells, and detection of anti-HTLV-1 antibodies in serum. ATL was subtyped according to JLSG criteria ^33^. The Comité de Protection des Personnes approved all experiments (DC-2019-3692). Supplemental Table S1 summarizes patient demographics and HTLV-1 proviral load data.

### NF-κB p65 Activation Assays

Nuclear extracts from primary CD8-depleted PBMCs of HTLV-1 asymptomatic carriers (AC) and patients with acute ATL (ATL) were prepared using the commercially available NE-PER™ Kit, according to the manufacturer’s instructions (Pierce). We then subjected these nuclear extracts to TransAM NF–κB p65 transcription factor ELISAs according to the manufacturer’s instructions (Active Motif). Briefly, 10 µg of protein extracts were allowed to bind to an oligonucleotide-coated plate. NF-kB–p65 was detected by incubation with specific primary antibodies and HRP-conjugated secondary Ab. The colorimetric readout was measured at 450 nm, with an optional reference wavelength of 655 nm, using a Spark 10M microplate reader (Tecan), and is proportional to transcription factor activity.

### RT-qPCR and Nanoplate-based digital PCR assay

Cells were collected and cryopreserved as dry pellets at-80°C until use. Nucleic acid was extracted using the AllPrep DNA/RNA Mini Kit (QIAGEN). 5 µg of total RNA isolated from each sample was subjected to reverse transcription using the All-In-One 5X RT MasterMix (abm) according to the manufacturer’s protocol to obtain first-strand cDNA. Transcript abundance was assessed by real-time quantitative (q)PCR using SYBR Green I Master Mix (Roche). Gene-specific primer sets used are listed in Supplementary Table S1. Standard curves for qPCR were generated for all primer pairs using serial dilution of appropriate experimental samples. We amplified samples in triplicate on each plate. We analyzed data using LightCycler®480 Software (Roche Diagnostics). Relative mRNA levels among experimental samples were determined using the 2−ΔΔCT method ^34^, with HPRT1 and GADPH as reference housekeeping genes. We performed nanoplate-based digital PCR (dPCR) using the same qPCR assay targeting Tax and HBZ on the QIAcuity One (QIAGEN) with the 8.5K 96-well Nanoplate and QIAcuity EG PCR Kit, according to the manufacturer’s instructions. The cycling conditions were the same as qPCR. We analyzed the data using the QIAcuity Software Suite. Primers used for qPCR are provided in Supplemental Table S2.

### PICO assay for quantification of protein interactions

HEK293T cells were transiently transfected with expression plasmids encoding Fra-2-HA, FLAG-JunD, FLAG-p65 (RELA), and HBZ-Myc using jetOPTIMUS® transfection reagent (Polyplus) according to the manufacturer’s instructions. Cells were harvested 48 hours after transfection, washed twice with ice-cold PBS supplemented with EDTA-free protease inhibitor cocktail, and lysed using the lysis buffer supplied in the PICO Amplification Core Kit (PICO BioScience GmbH). Protein-protein interactions were quantified using the Protein Interaction Coupling (PICO) assay, a digital PCR-based proximity assay enabling absolute quantification of protein complexes. Proteins of interest were detected using antibodies directed against HA, and FLAG epitopes conjugated to PICO labels according to the manufacturer’s recommendations. For absolute quantification experiments, labeled antibodies were used at saturating conditions, with a final concentration of 500 pM in the binding reaction. Equal volumes of cell lysate and antibody mixture were combined in 96-well PCR plates and incubated overnight at 4°C. Following incubation, samples were diluted according to the manufacturer’s protocol and mixed with QIAcuity Probe PCR Master Mix, Coupling dPCR Mix, and the corresponding PICO probe sets. We loaded reactions onto QIAcuity Nanoplates and analyzed them using a QIAcuity Digital PCR instrument (QIAGEN). We performed digital PCR using QIAcuity Nanoplate 8.5k 96-well plates. Thermocycling conditions consisted of an initial activation step at 95°C for 2 minutes, followed by 40 cycles of 95°C for 15 seconds and 58°C for 30 seconds. We acquired fluorescence signals using the FAM and HEX channels corresponding to the respective PICO probes. We analyzed digital PCR data using PIQuant software (PICO BioScience GmbH). Under saturating antibody conditions, interaction frequencies were determined by directly counting couplex-positive partitions and converted to absolute concentrations of protein complexes using Poisson statistics and partition-volume correction. We subtracted background signals measured in antibody-only controls from each sample before analysis. Results are reported as absolute numbers of protein complexes per microliter of lysate.

### Inflammatory Cytokines Enzyme-linked Immunosorbent Assay (ELISA)

Diluted serum samples from HTLV-1 asymptomatic carriers and patients with acute ATL were used for the Multi-Analyte ELISArray kit (MER-004A, QIAGEN) and the levels of various cytokines (IL-1α, IL-1β, IL-2, IL-4, IL-6, IL-10, IL-12, IL-17A, IFNγ, TNFα, GM-CSF) were determined from the absorbance read on a SPARK10 Microplate Reader (TECAN) at 450 and 570 nm according to the manufacturer’s protocol.

### Antibodies and plasmids

pcDNA3.1-HA-Fra-2 (OHu30302) was purchased from Genescript. pCDNA5-FRT-HA-cFos (human) was provided by the MGC platform (Biocampus-Montpellier). pcAG-Tax-Flag, pCMV-P65-Flag, pcDNA3.1 HBZ-Myc, and pcDNA3.1 JunD-Flag were previously described ^35,36^. N- and C-terminal deletions of Flag-P65 and HA-Fra-2 were PCR amplified from the HA-Fra-2 and Flag-P65 plasmids. Inserts were then cloned into the EcoR1-XhoI sites of the pGAGS-HA or the pGAGS-Flag plasmids, respectively. The primer sequences are provided in Supplementary Table S2. Anti-cFos (C-11) and Anti-JunD (329) were purchased from Santa Cruz Biotechnology. The anti-Fra-2 (HPA004817) was purchased from Sigma-Aldrich. The HTLV-1 Tax was detected using either hybridoma supernatant (AIDS reagent) or monoclonal anti-Flag M2 (Sigma-Aldrich). HBZ was detected using monoclonal anti-Myc (9E10) (Santa Cruz Biotechnology). NF-κB p65 was detected using the monoclonal anti-Flag M2 antibody (Sigma-Aldrich) or the L8F6 monoclonal antibody (Cell Signaling). Sp1 nuclear protein was detected using the Rabbit Polyclonal (ab12111) from Abcam. Actin and nucleolin were detected with anti-Actin (Sigma-Aldrich) and anti-C-23 (MS-3) (Santa Cruz Biotechnology), respectively. For the IL-1 neutralization assay, IL-1β-neutralizing antibody [4M27] was purchased from GENETEX, and Anakinra (AABH99A44F80) was purchased from Sigma-Aldrich.

### Cell Culture and Transfection

Human T-cell lines Jurkat (ATCC® TIB-152™), HuT-102 (ATCC® TIB-162™), and C8166 (ECACC 88051601) were propagated in RPMI-1640 with 10% fetal calf serum (FCS). ATL-2 (-) (CVCL_A6TF) were cultivated in ImmunoCult™-XF T Cell Expansion Medium (STEMCELL Technologies). HEK 293T cells (ATCC® CRL-3216™) were propagated in DMEM with 10% FCS and transfected according to the manufacturer’s protocol using PEIMax (1mg/ml, Polysciences). PBMCs were isolated from EDTA-anticoagulated blood samples from HTLV-1–infected patients without declared malignancy (AC) and Acute ATL patients (ATL) on Ficoll-density gradients and washed in phosphate-buffered saline (PBS). CD8^+^ cells were removed using anti-CD8 paramagnetic microbeads, following the manufacturer’s instructions (Miltenyi Biotec). CD8^+^-cell-depleted PBMC from ATL patients were then placed in low-binding culture wells (round-bottomed 24-well plate) at 10^6^/mL in 1 mL ImmunoCult™-XF T Cell Expansion Medium (STEMCELL Technologies) supplemented with 5 µL/mL of ImmunoCult™ Human CD3/CD28/CD2 T Cell Activator (STEMCELL Technologies) for 3 days, then only in ImmunoCult™-XF T Cell Expansion Medium.

### Luciferase Assay

Luciferase assays were performed as previously described ^37^ in a Spark 10M microplate luminometer (Tecan) with either the LightSwitch Assay Kit (Active Motif), the Genofax A and C kits (Yelen) or the Galacto-Star™ β-Galactosidase Reporter Gene Assay System (Applied Biosystem) according to the manufacturer’s instructions. The Firefly Luciferase or RenillaSP luciferase (RenSP) activities were normalized for transfection efficiency based on Renilla luciferase and the β-Galactosidase activities, respectively. Results were expressed as fold increases relative to the negative control (Mock). Each assay was conducted in triplicate.

### Immunoprecipitation analysis

For co-precipitation of transiently expressed proteins, HEK293T cells were transfected with expression vectors encoding the HA-Fra2 with an equimolar ratio of a vector encoding a Flag-epitope tagged NF-kB P65 protein using PEIMax transfection reagent (Polysciences). 48 hours after transfection, the cells were harvested using cell-removing buffer (40 mM Tris pH 7.5, 1 mM EDTA, and 150 mM NaCl) and pelleted by centrifugation. The cells were washed twice with cold PBS and resuspended in lysis buffer (50 mM HEPES, pH 7.5, 0.5 % Nonidet P-40, 150 mM NaCl, 2 mM EDTA, 20 mM β-glycerophosphate, and protease cocktail). Cell lysate corresponding to 50 µg of total protein was kept as an input. HA-tagged or Flag-tagged proteins were immunoprecipitated with EZview™ Red Anti-HA or anti-Flag M2 Affinity Gel (Sigma-Aldrich) for 16 hours at 4°C. In ATL-2 cells, Fra2 was immunoprecipitated by incubating the cell lysate with an anti-Fra2 antibody for 1hr at 4°C. Protein A/G Sepharose™ beads (Santacruz Biotech) were added overnight at 4°C. The beads/immune complexes were washed five times with 1 ml of the pre-chilled lysis buffer. The immune complexes were released from the beads by boiling in 1x Laemmli buffer (Bio-Rad). The whole-cell extracts and precipitated material were separated on 10% polyacrylamide gel and analyzed by western blotting.

### Cellular fractionation and Western Blot Analysis

Forty-eight hours after transfection, cells were harvested using cell resuspension buffer (CRB). Cytoplasmic (C) and Nuclear (N) extracts were prepared using the commercially available NE-PER™ Kit, according to the manufacturer’s instructions (Pierce). Total lysates were prepared in RIPA buffer, separated by electrophoresis on precast SDS-PAGE gels, and transferred to PVDF membranes (Bio-Rad) as previously described ^35^. Incubations with primary antibodies to detect Tax, HBZ, NF-kB P65, AP-1 transcription factor cFos, JunD, and Fra-2, Nucleoline C23 and α-actin were followed by incubations with the appropriate secondary antibodies conjugated with horseradish peroxidase (HRP) (GE Healthcare). After extensive washing, the protein bands were visualized using enhanced luminescence (Roche) and the ChemiDoc imaging system (Bio-Rad). Protein bands were quantified following scanning by ImageJ software^38^.

### Bimolecular Fluorescence Complementation (BiFC) analysis

The NF-κB p65 and Fra-2 coding regions were PCR-amplified from the Flag-P65 and HA-Fra-2 plasmids (GenScript), respectively. Inserts were then cloned into EcoR1-KpNI sites of the pBiFC-VN155(I152L) (Addgene #27097) and pBiFC-VC155 (Addgene #22011) that have the N-terminal (VN: 1-154) and C-terminal (VC: 155–238) moieties of Venus fluorescent protein. The primer sequences are provided in Supplementary Table S2. Forty-eight hours after transfection, the BiFC signal was visualized on a Nikon Eclipse Ts2-FL epifluorescence microscope and analyzed using a SPARK 10 microplate reader (TECAN).

### Chromatin Immunoprecipitation assay (ChIP)

Chromatin immunoprecipitation assays (ChIP) were performed as previously described ^39^. Briefly, proteins from 2 x 10^7^ ATL-2 cells were cross-linked to DNA with 1% formaldehyde in PBS for 10 minutes at room temperature. Nuclei were isolated by incubating cross-linked cells with 1 ml of Cell Fractionation Buffer Kit PARIS™ (Invitrogen) for 10 min on ice and then pelleted by centrifugation (500xg). The nuclei were washed with 500 μl of Cell Fractionation Buffer Kit PARIS™ (Invitrogen) and incubated with 500 µL of Nucleus Lysis Buffer for three hours on a rotator at 4°C. The chromatin was fragmented by incubating with 5 µl of Micrococcal Nuclease Solution (Thermo Scientific) at 37°C for 20 minutes, yielding an average size of 200-500 bp. The cell debris were pelleted by centrifugation (20,000×g, 4°C), and supernatants were collected. Chromatin was diluted in 50 mM Tris-HCl, 167 mM NaCl, 1 mM EDTA, 0.01% SDS, 1.1% Triton X-100, and protease inhibitor mix. To reduce nonspecific background, 5 µg of anti-Fra-2 (G-5)X (sc-166102X) or control IgG antibodies were preincubated with Dynabeads® Protein G (Invitrogen) according to the manufacturer’s instructions. Antibody-bead complexes were then added to chromatin samples and incubated overnight at 4°C. After incubation, the antibody-chromatin/bead complex was washed four times, and DNA was purified using DNA purification Spin Filters. Immunoprecipitated DNA was subjected to qPCR analysis with SYBR Green detection. ChIP-qPCR primers are shown in Table S3.

## Statistical analysis

Two-tailed paired Student’s t-test or one-way ANOVA test with Dunn’s multiple comparisons post-test using GraphPad PRISM 8 software. Data are presented as mean ± SD. Differences were considered not significant (ns) or significant at *=*p*<0.05, **=*p*<0.01, ***=*p*<0.001 and ****=*p*<0.0001.

## Results

### The Oncoprotein Fra-2 activates NF-kB p65 in ATL type I cells

To determine the molecular subtype of the primary ATL samples included in this study, we first quantified *tax* and *hbz* transcript expression using digital PCR (dPCR) (Fig. 1A-C). We selected this approach because Tax expression in ATL is typically extremely low, heterogeneous, and often restricted to a small fraction of leukemic cells, making accurate quantification particularly challenging. Using this highly sensitive approach, we quantified Tax mRNA in cultured primary CD8^+^-deprived peripheral blood mononuclear cells (PBMCs) isolated from HTLV-1-infected patients, asymptomatic (AC) or with Acute ATL (ATL), to establish their Tax expression status and classify each case according to the molecular framework proposed by Nosaka and Matsuoka ^5^. Next, we used a DNA-binding ELISA assay to assess NF-κB p65 activation in nuclear extracts from cultured primary CD8+-deprived PBMCs isolated from six AC and six ATL patients. We observed that NF-κB p65 binding activity is higher in ATL than in AC patients (**Figure 1D**, p<0,0001).

**Figure 1.**
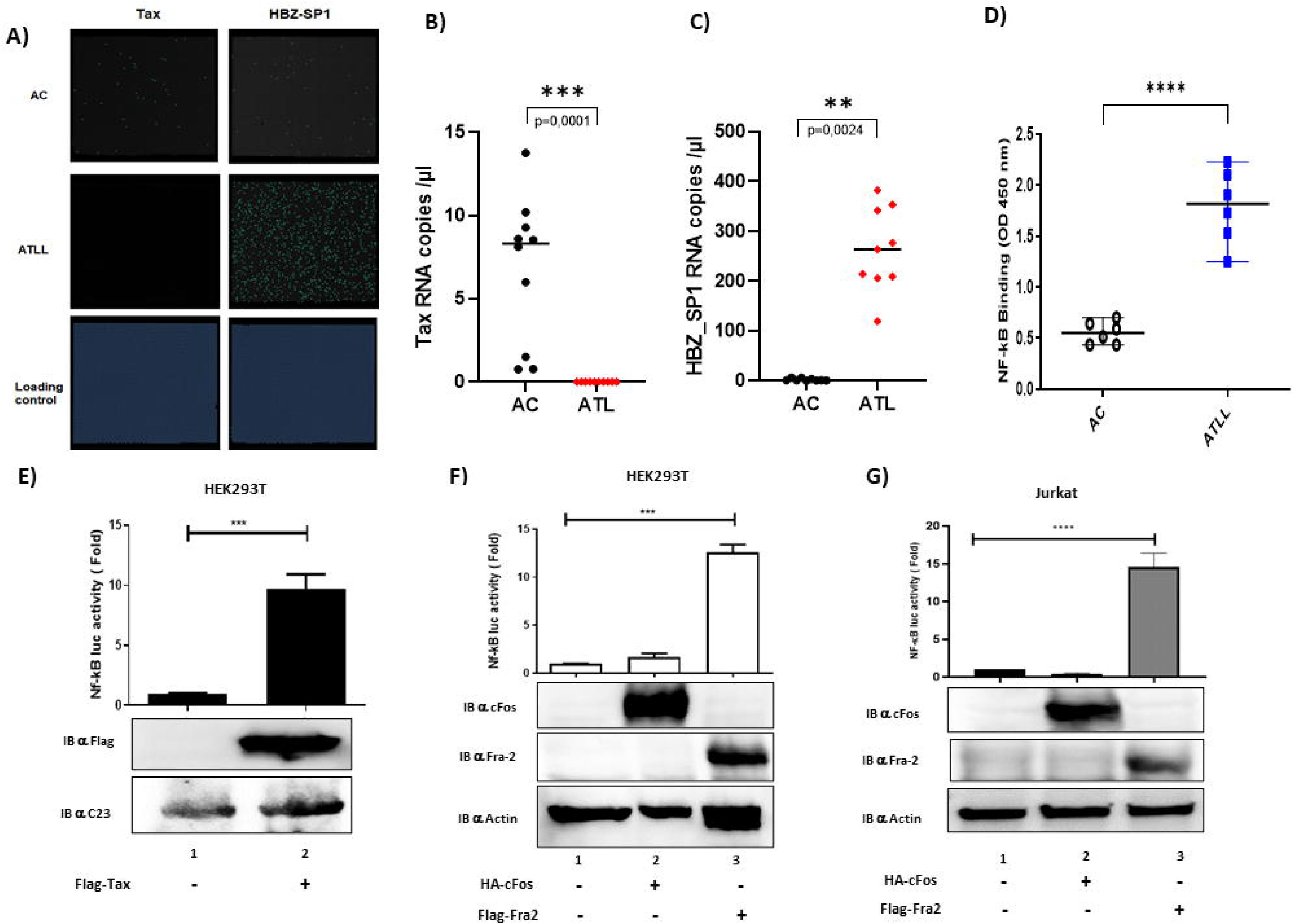
Fra-2 but not c-Fos enhances NF-κB transcriptional activity in T cells. (**A**) Digital PCR detection of Tax and HBZ_SP1 mRNA in CD8^+^-depleted PBMCs from HTLV-1 asymptomatic carriers (AC, n=6) and acute ATL patients (ATL, n=6). The lower panel shows the internal loading control. (**B-C**) Quantification of Tax and HBZ_SP1 transcript levels in AC and ATL patients using RT-qPCR and normalized with housekeeping gene. (**D**) NF-κB p65 DNA-binding activity in nuclear extracts from CD8^+^-depleted PBMCs (6 AC, 6 ATL) measured by TransAM NF-κB p65 ELISA (p < 0.0001, two-tailed Student’s t-test). (**E**) NF-κB reporter luciferase activity in HEK293T cells transfected with Flag-Tax (lane 2) or control vector (lane 1) and normalized to an internal control. Immunoblotting against the Flag tag confirms Tax expression.NF-κB reporter luciferase activity in (**F**) HEK293T and (**G**) Jurkat cells expressing HA-cFos or Flag-Fra-2. Representative immunoblots with respective antibodies confirm protein expression. Data are representative of 4 independent experiments. Statistical analysis: one-way ANOVA with Dunnett’s multiple comparisons test (*p < 0.01, **p < 0.001, ***p < 0.0001).

Several hypotheses remain under investigation to elucidate constitutive NF-κB activation in ATL cells. Some researchers believe constitutive NF-κB activation in ATL cells may result from extremely short bursts of Tax expression^40–42^, despite Zhao *et al.*’s report that the viral protein HBZ selectively suppresses the canonical NF-κB pathway by interacting with NF-κB p65 via its bZIP region ^43^. Understanding the mechanisms that regulate constitutive NF-κB activation in ATL cells is complex and essential to fully elucidate its role in intrinsic ATL chemoresistance. AP-1 transcription factors comprise proteins from the Jun, Fos, and ATF families and are activated by various stimuli, including stress and inflammatory cytokines. AP-1 regulates gene expression in cell proliferation, differentiation, and survival. Since Stein *et al.* have shown that NF-kB p65 can interact with the Fos and Jun transcription factors^44,45^, and Toualbi-Abed *et al.* have demonstrated that JunD cooperates with NF-kB p65 to activate the proximal NF-κB in the cyclin D1 promoter ^46^, we hypothesized that either a Fos, a Jun transcription factor, or another bZIP-containing protein could regulate NF-κB in ATL cells. To test whether cFos or Fra-2 influenced NF-κB activation and inflammatory cytokine transcription, we used a luciferase reporter assay to compare their ability to activate NF-κB in HEK293T and Jurkat T cells (**Figure 1E-G**). We observed that cFos did not increase transcription from the promoters (**Figures 1F and 1G, lane 2),** whereas Fra-2 significantly activated transcription from the NF-κB promoter, at the same level as our positive control Tax (Figure 1E), in both HEK293T and Jurkat cell lines (**Figures 1F and 1G, lane 3**).

### The Oncoprotein Fra-2 interacts with NF-kB p65

Following our recent observations, we next investigated potential interactions between Fra-2 and important NF-κB factors. Thus, we performed a Bimolecular Fluorescence Complementation Assay (BiFC) in HEK293T cells to test for interaction between Fra-2 and p65. In this assay, Fra-2 and p65 were respectively fused to the N- or C-terminal domains of the fluorescent Venus protein which will only fluoresce when both moieties are in direct interaction with each other (**Figure 2A, top panel**). After cotransfection of the recombinant plasmids into HEK293T cells, we assessed Venus fluorescence by epifluorescence microscopy. We observed that Venus fluorescence was restored in the nuclei of cells only when both pBiFC-VN155-p65 and pBiFC-VC155-Fra-2 were present, which was not the case when the p65 plasmid was cotransfected with the C-terminal Venus without Fra-2 fusion (**Figure 2A, bottom panel**). These results suggest that p65 and Fra-2 may be interacting in cells. To further confirm this interaction between Fra-2 and NF-κB p65, extracts from 293T or Jurkat cells transfected with Flag-tagged p65 and HA-tagged Fra-2 expression vectors were immunoprecipitated with either anti-Flag or anti-HA antibodies (**Figure 2B and 2C**), and an extract from an ATL patient-derived cell line, ATL-2, was immunoprecipitated with an anti-Fra-2 antibody and analyzed by Western blot (**Figure 2D**). Results show that Fra-2 is strongly associated with HEK293T or Jurkat cells’ exogenous p65 (**Figure 2B and 2C**) as well as ATL-2 cells’ endogenous p65 (**Figure 2D**). Next, we generated and used N-ter and C-ter deletion mutants of either NF-κB p65 or Fra-2 (**Figure 3A**) in order to narrow down the domain responsible for Fra-2-p65 interaction. Using a fluorescence complementation assay (BiFC) as described previously, we observed that Venus fluorescence is recovered only when the bZIP region of Fra-2 and the Rel domain of p65 are absent, meaning that those two domains are responsible for the interaction between Fra-2 and p65 (**Figures 3B and 3C**). These results were further confirmed by co-immunoprecipitation (**Figures 3D and 3E**).

**Figure 2.**
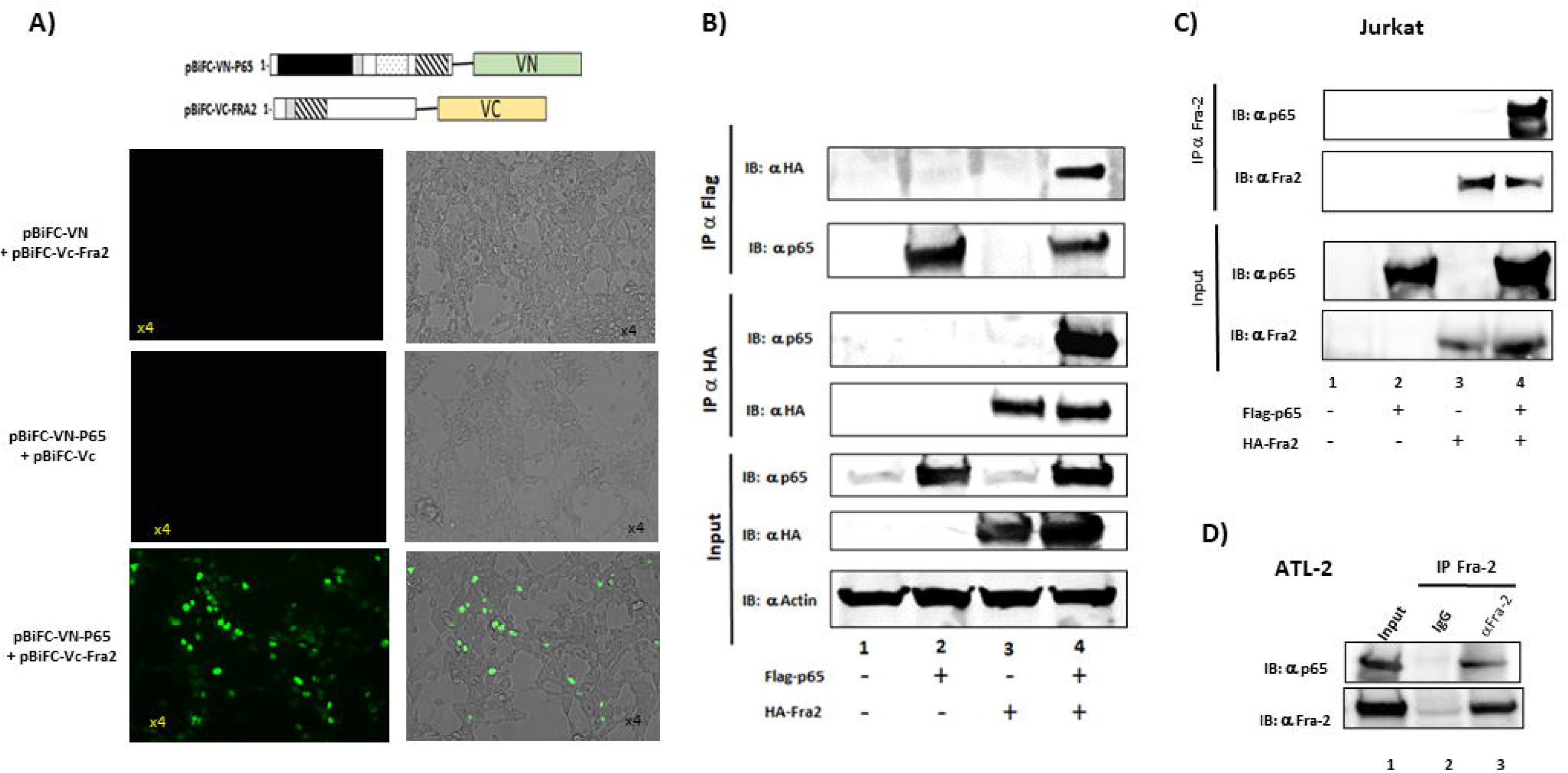
Fra-2 interacts with NF-κB p65. (**A**) Bimolecular fluorescence complementation (BiFC) assay using the p65 and Fra-2 constructs shown in the top panel, showing interaction between NF-κB p65 and Fra-2 in HEK293T cells. Venus fluorescence (green) indicates protein-protein interaction in cell nuclei. Co-immunoprecipitation of p65-Flag and Fra-2-HA overexpressed in (**B**) HEK293T cells and (**C**) Jurkat cells. 48 h after transfection, NF-κB p65 or Fra-2 was immunoprecipitated with anti-Flag or anti-HA magnetic beads, respectively, and interactions were visualized by immunoblotting against the other interactant. (**D**) Co-immunoprecipitation of endogenous Fra-2 and p65 in ATL-2 cells. Fra-2 was immunoprecipitated with anti-Fra-2 antibody, and bound p65 was detected by immunoblotting.

**Figure 3.**
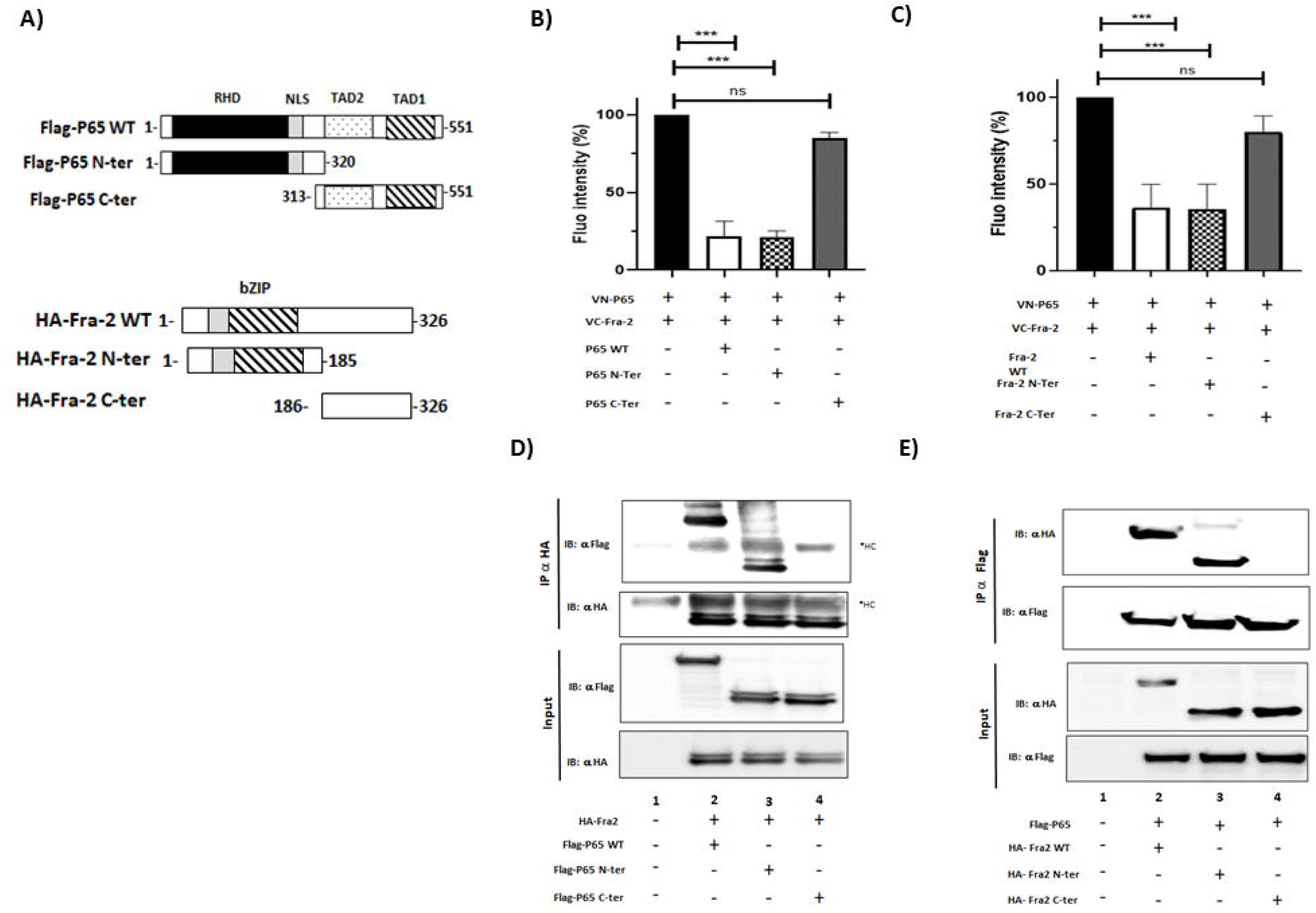
Mapping of Fra-2 and p65 interaction domains. (**A**) Schematic representation of NF-κB p65 and Fra-2 truncation mutants used in this study. Rel homology domain (RHD) of p65 and bZIP domain of Fra-2 are indicated. (**B-C**) Quantification of BiFC assay between Fra-2 WT and p65 truncation mutants. HEK293T cells were co-transfected with pBiFC-VC-Fra-2, pBiFC-VN-p65, the p65 mutants, or the Fra-2 mutants. 48 h after transfection, Venus fluorescence was measured using a Spark 10M microplate reader (TECAN) (excitation 514 nm, emission 529 nm). (**D–E**) Co-immunoprecipitation assays confirming domain requirements for the Fra-2/p65 interaction. Data are representative of 4 independent experiments. Statistical analysis: one-way ANOVA with Dunnett’s multiple comparisons test (*p < 0.01, **p < 0.001, ***p < 0.0001).

Upon stimulation by proinflammatory cytokines (e.g., IL-1α or IL-1β, TNFα), the cytosolic NF-κB pathway generates active DNA-binding dimers that translocate to the nucleus and bind available NF-κB binding sites, thereby reprogramming the expression of numerous genes^17,47,48^. To investigate how Fra-2 could potentially activate NF-κB, we next used an NF-κB reporter assay (p3κb-Luc). In short, HEK293T and Jurkat cells were cotransfected with either cFos or Fra-2 along with p65 (**Figures 4A and 4B**). While cotransfection of cFos or HTLV-1 Tax did not affect the activation of NF-κB by p65 (**Figures 4C and 4D),** Fra-2 significantly activates it, as shown by both luciferase activity and protein level increases (**Figures 4A and 4B**). Next, we performed cellular fractionation to determine whether Fra-2 increases NF-κB activation by facilitating p65 nuclear translocation. We observed that, with or without Fra-2 expression, a similar amount of endogenous p65 was found in the nuclear fraction (**Figure 4F, lines 2 and 4**), which means that Fra-2 doesn’t impact p65 nuclear localization. Thus, we hypothesized that binding to Fra-2 could instead alter NF-κB p65 DNA-binding activity. To test this, we used a DNA-binding ELISA assay to probe for NF-κB p65 activation in nuclear extracts from HEK293T cells transfected with an empty vector (Mock), p65-Flag, or p65-Flag and Fra-2-HA. Using a similar amount of p65-Flag (**Figure 4G),** we observed higher p65 DNA-binding activity when Fra-2 was co-expressed **(Figure 4H, white vs. gray histogram, p = 0.0006**), indicating that Fra-2 binding to p65 activates NF-κB via p65

**Figure 4.**
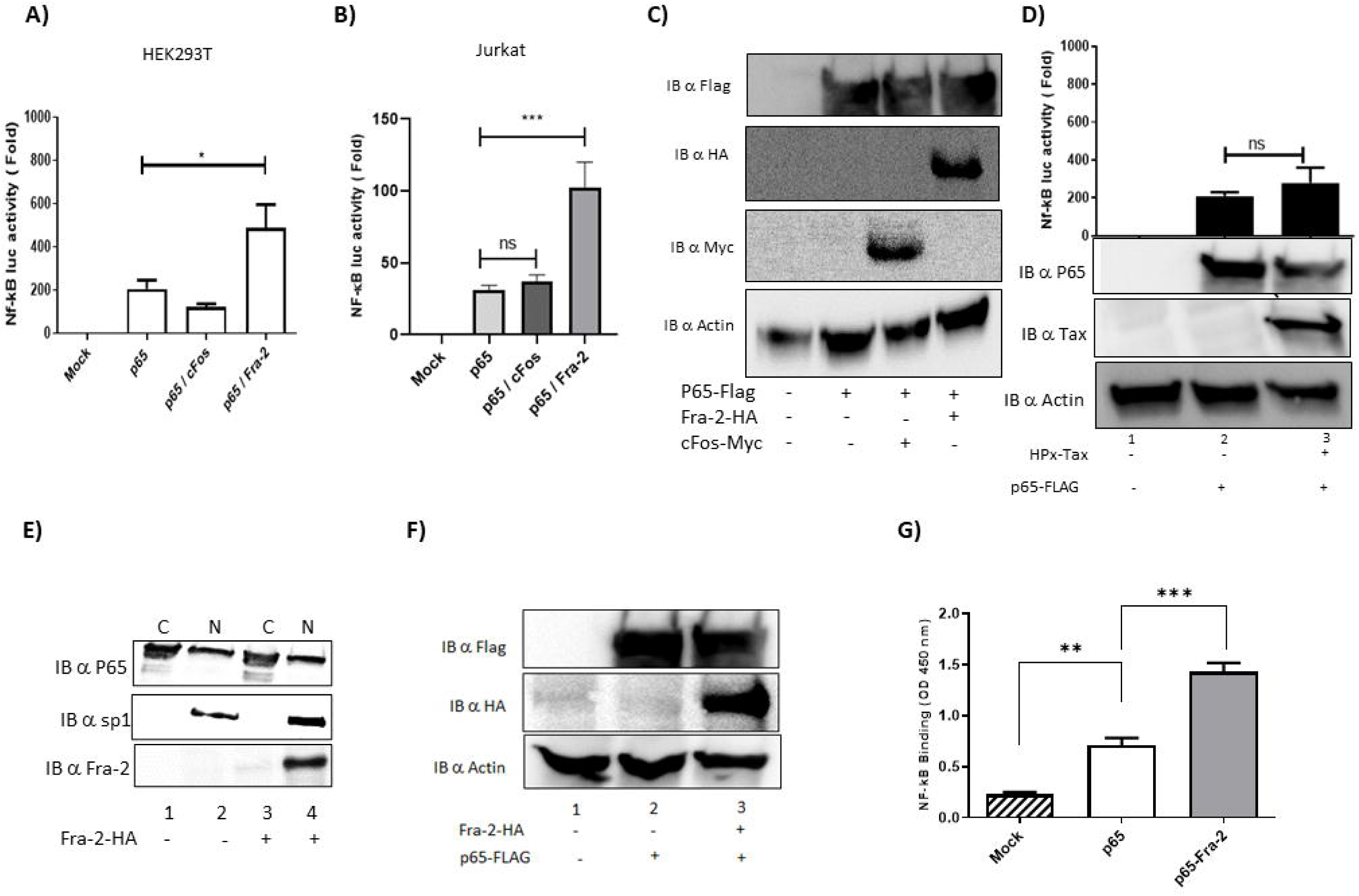
Fra-2 enhances NF-κB p65 DNA-binding activity. NF-κB reporter luciferase activity in (**A**) HEK293T and (**B**) Jurkat cells expressing p65 along with c-Fos or Fra-2. Cells were co-transfected with an NF-κB-luciferase reporter plasmid and the indicated proteins. 48 hours after transfection, luciferase activity was measured and normalized to an internal control. (**C**) Immunoblot confirming protein expression in Jurkat cells. (**D**) NF-κB reporter activity in the presence of HTLV-1 Tax. HEK293T cells were co-transfected with an NF-κB-luciferase reporter plasmid, p65, and with or without Tax. (**E**) Immunoblot of cytoplasmic (C) and nuclear (N) fractions showing p65 localization in the presence or absence of Fra-2. Sp1 served as a nuclear-fraction control. (**F**) Immunoblot confirming Fra-2 and p65 expression in total cell lysates. (**G**) NF-κB p65 DNA-binding activity in HEK293T cells transfected with Flag-p65 alone or with HA-Fra-2, measured using the TransAM NF-κB p65 ELISA. Data are representative of 4 independent experiments. Statistical analysis: one-way ANOVA with Dunnett’s multiple comparisons test (*p < 0.01, **p < 0.001).

### HBZ increases NF-κB activation mediated by Fra-2

Fra-2 is part of the AP-1, another critical transcription factors family that is known to cooperate with NF-κB to further enhance chemoresistance in ATL^27^. In ATL, AP-1 activation contributes to chemoresistance by promoting gene expression that supports cell survival and proliferation and by inhibiting apoptotic pathways. Because HTLV-1 HBZ interacts with multiple transcriptional regulators including AP-1 factors, we next examined its effect on Fra-2– dependent NF-κB activation. Expression of HBZ significantly increased NF-κB reporter activity in both HEK293 (data not shown) and Jurkat cells, indicating that HBZ positively modulates transcriptional activity mediated by Fra-2 and p65 interaction, which is not the case for p65 alone or p65 withJunD, another AP-1 factor (**Figure 5A**). Subsequent co-immunoprecipitation experiments showed that Fra-2 interacted with JunD and that this interaction was reduced by half in the presence of HBZ, suggesting that HBZ interferes with AP-1–related transcriptional complexes (**Figure 5B, lanes 4 and 8**). In contrast, HBZ enhanced or stabilized the interaction between Fra-2 and p65, as reflected by increased co-immunoprecipitation efficiency (**Figure 5C, lanes 4 and 8**).

**Figure 5.** HBZ modulates Fra-2–dependent NF-κB transcriptional activity. (**A**) NF-κB reporter luciferase activity in Jurkat cells expressing p65 and Fra-2 in the presence or absence of HBZ. Cells were co-transfected with an NF-κB-luciferase reporter plasmid, p65, JunD, and Fra-2 alone or in a dimer with p65, and in the presence or absence of HBZ-Myc. We measured luciferase activity 48 hours after transfection. Immunoblotting confirms expression of p65, Fra-2, JunD, and HBZ. Co-immunoprecipitation assays showing (**B**) Fra-2-HA/JunD-Flag and (**C**) Fra-2-HA/p65-Flag interactions in the presence or absence of HBZ-Myc. HEK293T cells were co-transfected with p65-Flag, JunD-Flag, and Fra-2-HA alone or with HBZ-Myc. 48 hours after transfection, JunD-Flag (**B**) or Fra-2-HA (**C**) was immunoprecipitated, and bound proteins were detected by immunoblotting. We quantified band intensities using ImageJ software and normalized to the corresponding actin band intensity. (**D**) 2D-multiplex plot showing the Fra-2-HA/JunD-Flag complex in the absence or presence of HBZ-Myc as analyzed by PICO assay and quantified in (**E**) using PIQuant software (p < 0.001, Student’s t-test). (**F-G**) 2D-multiplex plot and associated quantification showing the Fra-2-HA/p65-Flag complex in the presence or absence of HBZ. (p < 0.001, Student’s t-test). Data are representative of 3 independent experiments.

Next, to verify the protein interactions and the impact of HBZ on the complexes, we employed the Protein Interaction Coupling (PICO) assay, a digital immunoassay that blends antibody-based target detection with compartmentalized amplification and sensing on the QIAcuity Digital PCR platform. Briefly, 48 hours after transfecting cells with Fra-2-HA/JunD-Flag or Fra-2-HA/P65-Flag, with or without HBZ-Myc, we incubated cell lysates overnight with oligonucleotide-labeled antibodies targeting HA or Flag tags. This step allowed proximity-dependent antibody complexes to form when interacting proteins were present. After dilution, we transferred the samples to QIAcuity 8.5k 96-well Nanoplates, where each was divided into around 8,500 pico-liter reaction chambers. Inside these chambers, the coupled antibody oligonucleotides were amplified and detected through multiplex digital PCR (**Fig. 5D and 5F**). We then measured the fluorescent signals from individual labels and complexes across all the partitions, enabling highly sensitive detection of protein interactions with very low background noise. To quantify these interactions, we used Poisson statistics with PIQuant software, which gave us a clear measure of how often proteins connected and their interaction ratios. We observed that the Fra-2/JunD complex was decreased by 8-fold in the presence of HBZ (**Fig. 5E),** while the Fra-2/P65 dimer was increased by 5.3-fold by HBZ (**Fig. 5G**). Together, these data demonstrate that HBZ differentially regulates Fra-2–containing transcriptional complexes, favoring NF-κB–associated interactions such as p65, over classical AP-1 complexes like JunD.

### The Oncoprotein Fra-2 increases DNA binding by NF-κB p65 and activates the promoters of inflammatory cytokines

Cytokines are critical signaling proteins produced within the cancer microenvironment that significantly influence tumor progression and treatment response. To better understand chemoresistance in ATL patients, we compared the expression levels of 12 inflammatory cytokines in the HTLV-1-derived cell lines Hut102, C81-66, and ATL-2 (**Figure 6A**) and in cultured primary CD8^+^-deprived PBMCs isolated from AC and ATL patients by qPCR (**Figure 6B**). IL-1α, IL-1β, IL-2, IL-17α, TNFα, and GM-CSF transcript levels were higher in the ATL-2 cell line than in the Jurkat cell line (**Figure 6A**). Similar observations were made in the ATL patient cells when compared to ACs (p = 0.0001) (**Figure 6B**), indicating a consistent overexpression of inflammatory cytokines in ATL. Next, using a Multi-Analyte ELISArray kit, we compared the expression levels of twelve inflammatory cytokines (IL-1α, IL-1β, IL-2, IL-4, IL-6, IL-8, IL-10, IL-12, IL-17, IFNγ, TNFα, and GM-CSF) in sera from Caribbean HTLV-1 asymptomatic carriers and ATL type I patients. We observed that sera from ATL type I patients were enriched for IL-1α, IL-1β, IL-2, IL-10, IL-12, IL-17α, TNF-α, and GM-CSF (**Figure 6C**), further strengthening our previous results. Next, we performed a correlation analysis to assess a potential correlation between cytokine mRNA levels and proviral load (PVL) in HTLV-1-infected individuals (**Figure S2**). No significant correlation was highlighted between the HTLV-1 PVL and the upregulation of cytokine transcripts.

**Figure 6.**
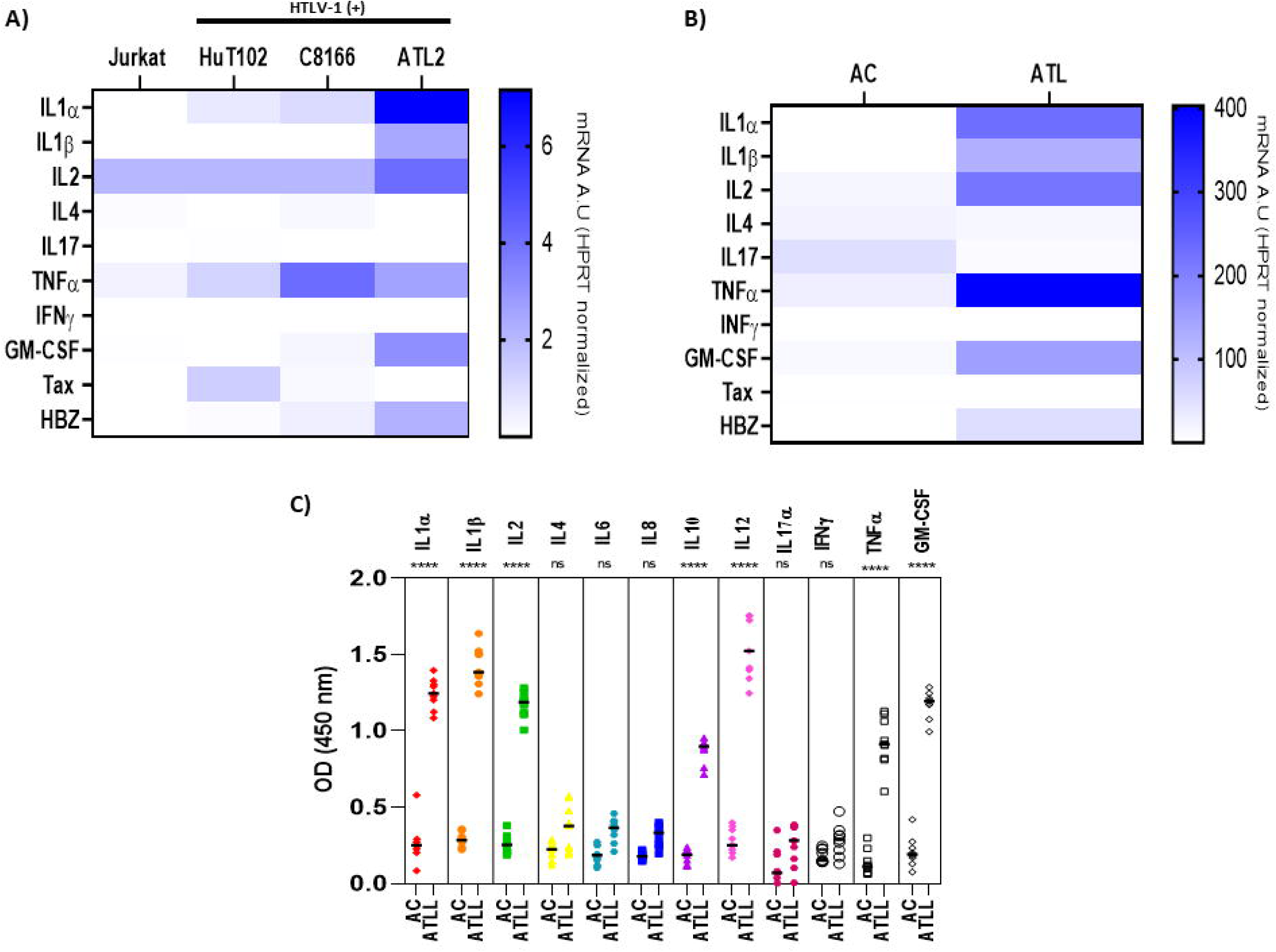
Increased inflammatory cytokine expression in Type I ATL patients. (**A**) Heatmap of relative mRNA expression of cytokines (IL1A, IL1B, IL2, IL4, IL17A, CSF2, TNF) and viral transcripts (Tax, HBZ) in Jurkat, HuT-102, C8166, and ATL-2 cells, measured by RT-qPCR. (**B**) Heatmap of relative mRNA expression of the same cytokines in CD8^+^-depleted PBMCs from HTLV-1 asymptomatic carriers (AC, n=6) and acute ATL patients (ATL, n=6). (**C**) Cytokine secretion measured by Multi-Analyte ELISArray Kit in serum from asymptomatic carriers (AC, n=6) and Type I ATL patients (ATL, n=6). Optical density (OD 450 nm, reference: 570 nm) is shown for IL-1α, IL-1β, IL-2, IL-4, IL-6, IL-8, IL-10, IL-12, IL-17A, IFN-γ, TNF-α, and GM-CSF. Each symbol represents an individual sample; bars indicate mean ± SEM. Data are representative of 4 independent experiments. Statistical analysis: one-way ANOVA (*p < 0.01, **p < 0.001).

Finally, we investigated the role of Fra-2 in activating proinflammatory cytokines expressed in ATL cells. For that, we cotransfected HEK293T cells with the promoters of Il-1β, Il-2, Il-4, IL-17, and CSF2 (GM-CSF) and either an empty vector (Mock), Flag-P65, HA-Fra2, or HTLV-1 Tax (**Figures 7 and S5)**. We collected cells two days after transfection and measured luciferase activity. While p65 is seen to activate the promoters of various cytokines, we observed that co-expression of p65/Fra-2 significantly increases activation of the promoters of Il-1β (p<0,0001), IL-2 (p<0,0001), and CSF2 (p<0,0001) but not the promoters of Il-4 (p=0,7079) and IL-17 (p=0,9996). Fra-2 alone did not affect Il-4 or Il-17 **(Figures 7A-7E).** We also showed that Tax only weakly activated the transcription of Il-1β and CSF2 (**Figure S5**) compared to Fra-2 alone or the cotransfection Fra-2/p65 dimer **(Figures 7A-7E).** Together, these observations suggest that Fra-2-mediated activation of NF-κB is responsible, at least in part, for the overexpression of pro-inflammatory cytokines as observed in ATL.

**Figure 7.**
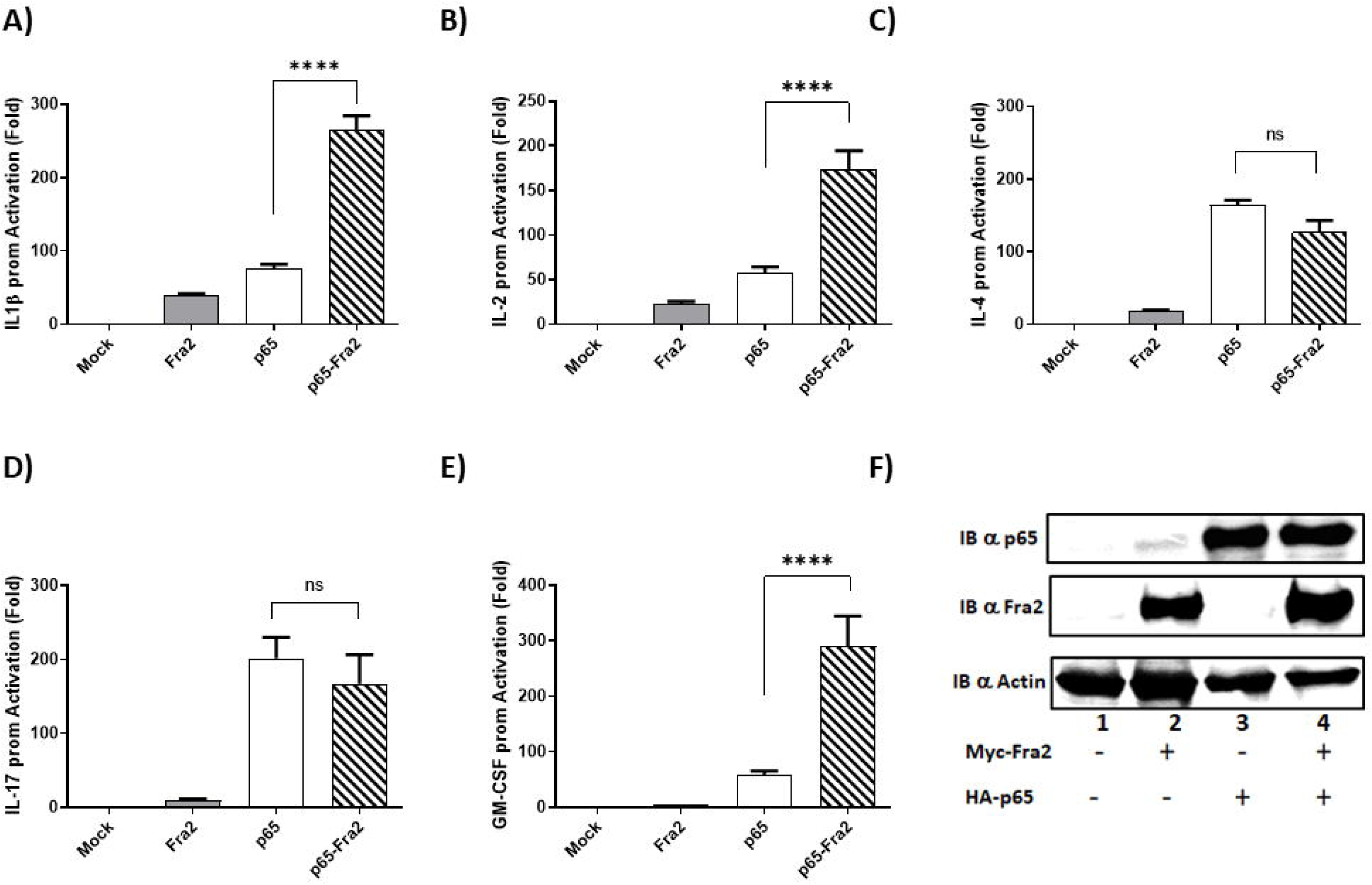
Fra-2 and p65 cooperatively activate pro-inflammatory cytokine promoters. Promoter activity of (**A**) IL-1β, (**B**) IL-2, (**C**) IL-4, (**D**) IL-17, and (**E**) CSF2 (GM-CSF) measured by luciferase reporter assay. HEK293T cells were co-transfected with reporter luciferase promoter plasmids for the indicated cytokines along with HA-p65-Flag, Myc-Fra-2, or Mock. 48 hours after transfection, luciferase activity was measured. (**F**) Immunoblot confirming p65 and Fra-2 expression; actin served as a loading control. Data are representative of 4 independent experiments. Statistical analysis: one-way ANOVA with Dunnett’s multiple comparisons test (ns p > 0.05, *p < 0.01, **p < 0.001, ***p < 0.0001).

### HBZ and Fra-2 promote IL-1β transcription, and IL-1 signaling contributes to ATL cell survival

Because Fra-2 and p65 strongly induced IL-1β, we investigated HBZ’s effect on IL-1β transcription. Co-expression of HBZ with Fra-2 and p65 further increased IL-1β promoter activation mediated by Fra-2 and p65 (Figure 8A, white histogram vs Black histogram), suggesting an activating effect of HBZ. Immunoblot analysis confirmed that HBZ did not affect Fra-2 or p65 expression (**Figure 8B**). To further analyze the recruitment of Fra-2 on the promoters of inflammatory cytokines *in cellulo* within the physiologically relevant context of the chromatin structure, we first identified the NF-kB and AP-1 binding sites throughout the promoters of IL-1β, IL-2, TNF-α and GM-CSF using Tfsitescan and PROMO ^49^ (**Figure S6**). Next, we performed chromatin immunoprecipitation (ChIP) experiments on nuclear extracts from ATL-2, since this ATL-derived cell line expresses high levels of Fra-2 and IL-1β, Il-2, TNF-α, and GM-CSF mRNA (**Figure 6**). ChIP with an anti-Fra-2 antibody showed that Fra-2 occupancy is primarily at NF-κB binding sites and not at the AP-1 binding sites in the promoters of IL-1β (**Figure 8C**), IL-2, TNF-α, and GM-CSF (**Figure S6**). These observations highlight the role of the Fra-2/p65 complex in activating inflammatory cytokines in ATL cells. To assess the functional relevance of IL-1β signaling in the chemoresistance observed in ATL cells, ATL-2 cells or CD8^+^-depleted PBMCs from ATL type I patients were pretreated for four hours with either the IL-1 receptor antagonist Anakinra (**Figure 8D-8E**) or a neutralizing anti–IL-1β antibody (**Figure 8F-8G**) and then submitted to increasing concentrations of doxorubicin for 24 hours. We observed that IL-1 inhibition increased sensitivity to doxorubicin and significantly reduced ATL-2 cell or CD8-depleted PBMCs from ATL type I patients’ survival, with IC50 values of 1.6 µM for Anakinra and 5.2 µM for the neutralizing anti-IL-1β, respectively (**Figure 8D–G**). These findings indicate that Fra-2 and HBZ cooperate to promote IL-1β expression and that IL-1β signaling in return contributes to the chemoresistance of ATL type I cells.

**Figure 8.**
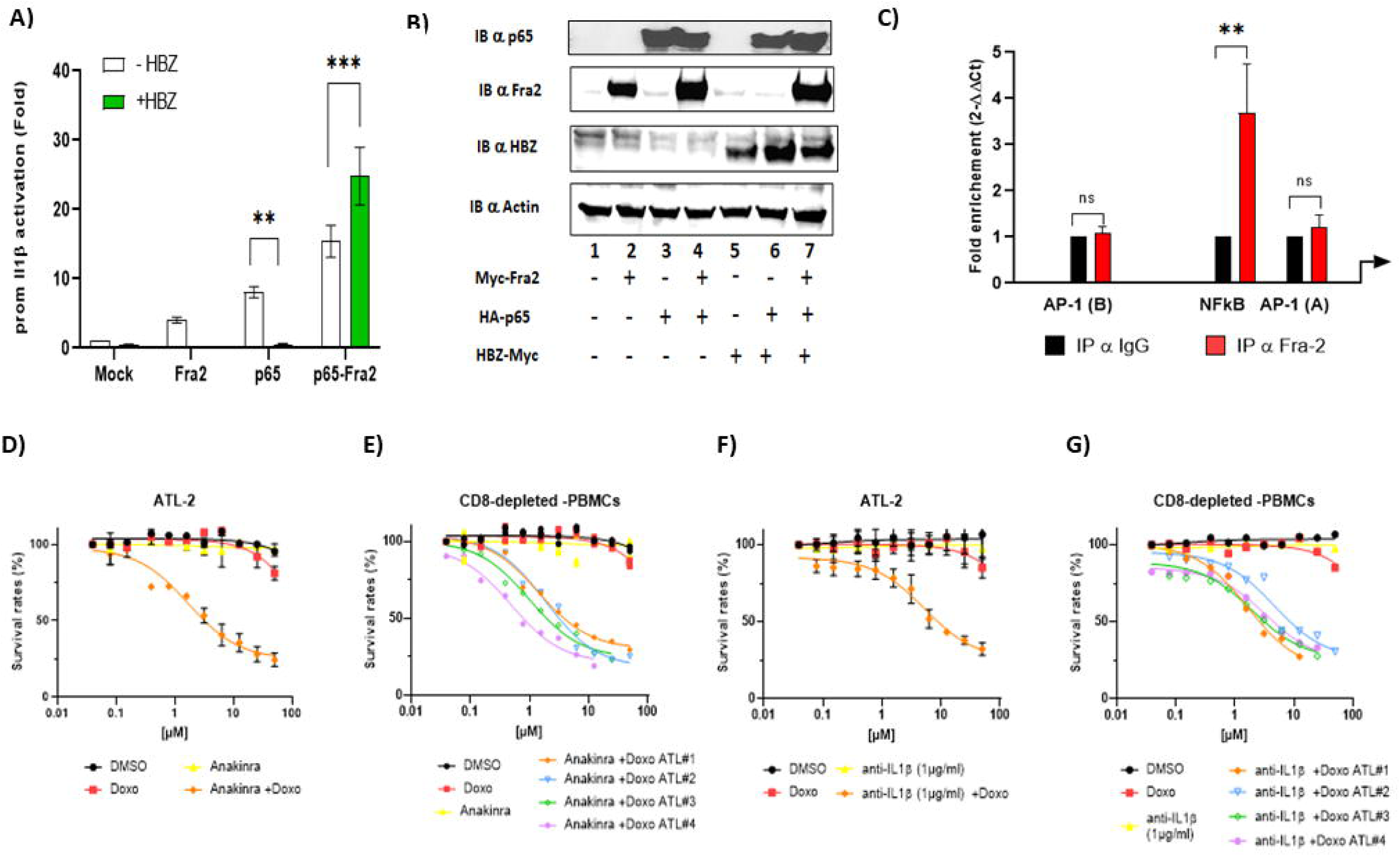
HBZ enhances IL-1β transcription, and IL-1 signaling contributes to ATL cell survival. (**A**) IL-1β promoter activity in HEK293T cells expressing Fra-2, p65, or both in the presence or absence of HBZ.. 48 hours after transfection, luciferase activity was measured (p < 0.0001, one-way ANOVA with Dunnett’s test). (**B**) Immunoblot confirming HBZ, p65, and Fra-2 expression; actin served as a loading control. (**C**) Chromatin immunoprecipitation (ChIP) showing Fra-2 enrichment at NF-κB and AP-1 binding sites within the IL1B, IL2, TNF, and CSF2 promoters in ATL-2 cells. Data are presented as fold enrichment relative to IgG control (*p < 0.01, **p < 0.001, two-way ANOVA). (**D–E**) ATL-2 cell viability after 4 hours of pre-treatment with Anakinra (10 µg/mL) followed by doxorubicin treatment for 24 hours. Cell viability was assessed using PrestoBlue vital dye and survival rate was calculated relative to DMSO controls. (**F–G**) Viability of CD8^+^-depleted PBMCs from Type I ATL patients (n=4) after 4 hours of pre-treatment with anti-IL-1β antibody (clone 4M27, 1 µg/mL) followed by doxorubicin treatment for 24 hours. IC₅₀ values: 1.6 µM (Anakinra), 5.2 µM (anti-IL-1β) (p < 0.001, one-way ANOVA). Data are representative of 4 independent experiments.

**Figure 9.**
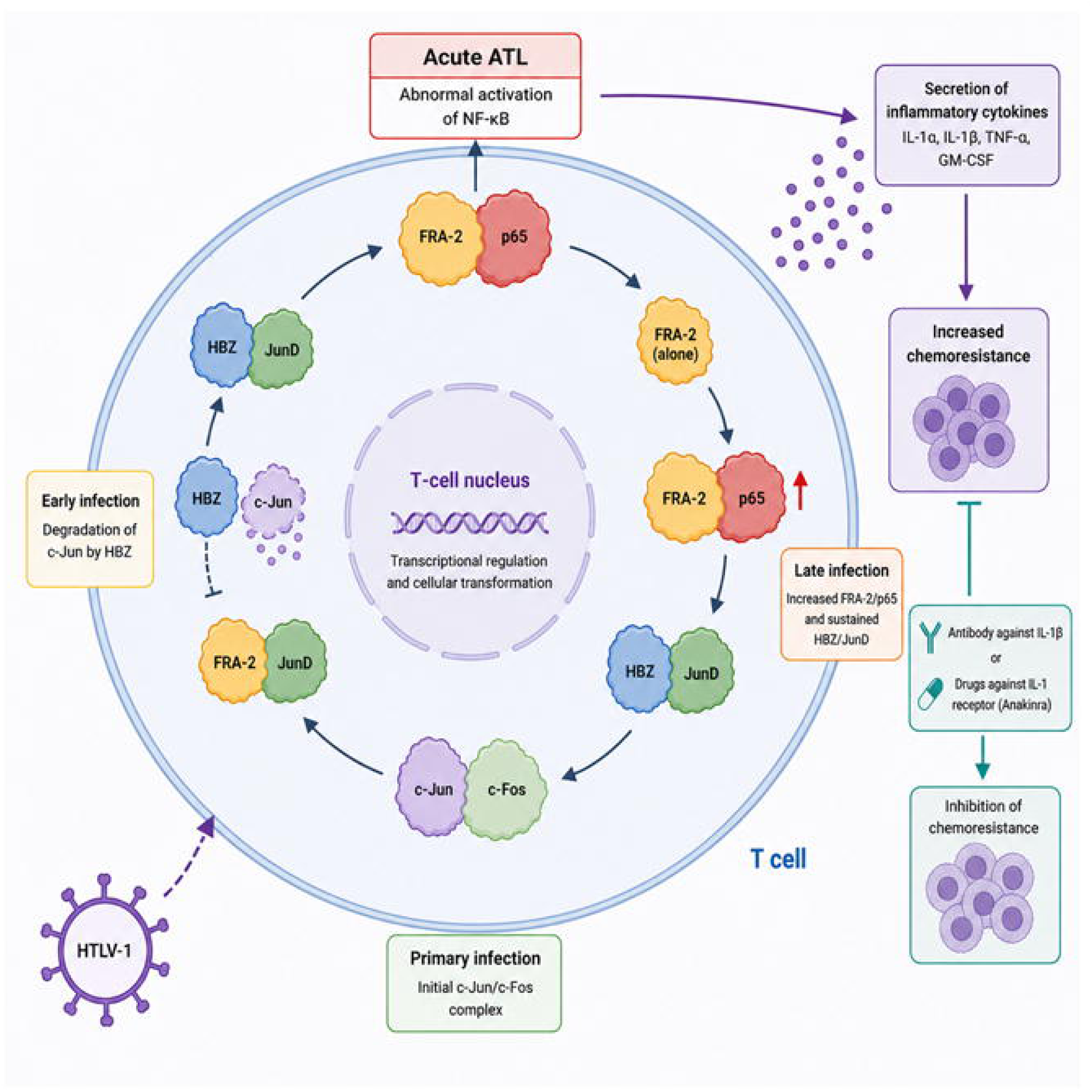
Graphical Abstract. Proposed model: HBZ reshapes Fra-2–containing transcriptional complexes to favor cooperation with NF-κB, resulting in enhanced IL-1β expression that sustains inflammatory signaling, promotes ATL cell survival, and contributes to chemoresistance.

## Discussion

Adult T-cell leukemia/lymphoma (ATL) is an aggressive malignancy caused by human T-cell leukemia virus type 1 (HTLV-1) infection and is associated with poor responses to chemotherapy and high relapse rates ^1–4^. Increasing evidence indicates that constitutive inflammatory signaling contributes not only to leukemogenesis but also to therapeutic resistance ^17–19^. In this study, we identify Fra-2 as a previously unrecognized regulator of inflammatory transcriptional programs in HTLV-1–infected cells and demonstrate that Fra-2 cooperates with NF-κB to promote IL-1β expression, thereby enhancing ATL type I cell survival and chemoresistance. We further show that the viral protein HBZ modulates Fra-2– containing transcriptional complexes and reinforces this inflammatory circuit, providing a mechanistic link between viral gene expression, dysregulated cytokine production, and resistance to cytotoxic therapy.

NF-κB signaling, particularly through the p65 (RelA) subunit, is a central regulator of genes involved in inflammation, cell survival, and proliferation ^48^. Constitutive activation of NF-κB in ATL type I has been linked to increased expression of anti-apoptotic proteins, cell-cycle regulators, and cytokines that promote leukemic cell survival ^20,28,50^. Our results extend these observations by identifying Fra-2 as a functional partner of NF-κB in this context. Although Fra-2 is traditionally considered a component of AP-1 complexes ^51^, our data demonstrate that Fra-2 enhances NF-κB transcriptional activity, promotes nuclear accumulation and DNA binding of p65, and selectively augments transcription of promoters for inflammatory cytokines. These findings indicate that Fra-2 can act outside classical AP-1 signaling to amplify NF-κB–dependent transcriptional programs.

Fra-2 has been implicated in regulating genes involved in cell proliferation, stress adaptation, and survival, including cyclin D1 and matrix metalloproteinases ^52–55^. Its overexpression has been associated with aggressive behavior and therapy resistance in several malignancies ^39,56–59^. The ability of Fra-2 to cooperate with NF-κB may therefore represent a mechanism by which leukemic cells integrate stress responses with inflammatory signaling to survive cytotoxic insults. Our findings also identify HBZ as a modulator of Fra-2–dependent transcriptional networks. HBZ altered protein–protein interactions involving Fra-2, reducing association with AP-1 partners while favoring interaction with p65. This shift in transcriptional complex composition suggests that HBZ may redirect transcriptional programs toward NF-κB– dependent inflammatory pathways, thereby reinforcing survival signals in ATL cells. These observations extend previous studies demonstrating a central role for HBZ in maintaining the proliferation and persistence of HTLV-1–transformed cells ^27,60–63^.

Cytokines play a central role in shaping the tumor microenvironment and influencing tumor progression and therapeutic response^17–19^. These signaling molecules regulate immune responses, angiogenesis, and survival pathways, and in hematologic malignancies, they can create a permissive niche that supports leukemic cell persistence ^17–19^. Our analysis of CD8-depleted PBMCs from asymptomatic carriers and ATL patients demonstrated increased production of inflammatory cytokines and enhanced NF-κB activity in ATL samples, consistent with previous observations that chronic inflammatory signaling is a hallmark of HTLV-1– associated leukemogenesis ^3^. These findings support a model in which dysregulated cytokine production contributes to both disease maintenance and treatment resistance.

A key downstream mediator identified in this study is IL-1β. IL-1 is a pro-inflammatory cytokine that plays a central role in immune regulation and tumor-associated inflammation^22,32,64,65^. IL-1 signaling promotes a pro-survival microenvironment by activating NF-κB and AP-1 pathways, inducing secondary cytokine production, and establishing positive feedback loops that sustain inflammatory signaling ^22,32,64,65^. Increasing evidence indicates that IL-1β contributes to chemoresistance in multiple malignancies by enhancing survival signaling and protecting tumor cells from apoptosis ^66–68^. Consistent with these observations, inhibition of IL-1 signaling in our study sensitized ATL cells to doxorubicin, supporting a functional role for IL-1β in therapeutic resistance and suggesting that inflammatory cytokine networks actively contribute to treatment failure in ATL.

These findings have potential therapeutic implications. Current treatment strategies for aggressive ATL often rely on CHOP-based chemotherapy, but responses are often incomplete, and relapse is common ^69–71^. Notably, our data demonstrate that Anakinra restores doxorubicin sensitivity in primary ATL cells (Figures 8D–E), providing preclinical evidence for this approach. Targeting pathways that sustain leukemic cell survival may improve treatment efficacy. For example, inhibition of NF-κB signaling with proteasome inhibitors such as bortezomib has been shown to enhance chemosensitivity in preclinical models ^72^. Similarly, targeting AP-1 activity or Fra-2–dependent transcription may reduce stress-adaptation pathways that contribute to resistance. Our results further suggest that blockade of IL-1 signaling, for example, with the IL-1 receptor antagonist anakinra ^73,74^, could disrupt inflammatory feedback loops that support leukemic cell survival. More broadly, our findings suggest that immunomodulatory approaches administered prior to chemotherapy may improve outcomes in ATL. Immunotherapy, or targeted inhibition of inflammatory pathways, before CHOP treatment may reduce survival signals, alter the tumor microenvironment, and increase leukemic cell susceptibility to cytotoxic agents. Antibody-based and immune-modulating strategies have demonstrated promising activity in ATL ^75,76^, and sequential or combination approaches that incorporate inhibition of cytokine pathways may warrant further clinical investigation.

In summary, we identify a Fra-2/NF-κB/IL-1β axis that links viral transcriptional regulation to inflammatory signaling and chemoresistance in ATL patients. Fra-2 cooperates with NF-κB p65 to drive cytokine expression, while HBZ reinforces this transcriptional program, promoting leukemic cell survival under therapeutic stress. These results support a model in which inflammatory transcriptional circuits actively contribute to treatment resistance rather than representing a secondary consequence of transformation. However, the precise molecular mechanisms by which Fra-2 promotes nuclear retention of p65 and how HBZ selectively reshapes transcriptional complexes remain to be elucidated. Targeting components of this pathway, including IL-1 signaling or upstream regulators of NF-κB activity, may therefore represent a rational strategy to enhance chemosensitivity and improve outcomes in patients with ATL.

### Graphical model statement

We propose a model in which HBZ reshapes Fra-2–containing transcriptional complexes to favor cooperation with NF-κB, resulting in enhanced IL-1β expression that sustains inflammatory signaling, promotes ATL cell survival, and contributes to chemoresistance.

## SUPPLEMENTAL FIGURES

**Supplemental Figure S1. Overview of nanoplate dPCR protein coupling assay (PICO).** Workflow schematic of the Protein Interaction Coupling (PICO) assay for quantification of protein-protein interactions. (1) Target-specific antibodies are conjugated with unique DNA oligonucleotides (PICOglue Labels) to form ready-to-use binding agents. (2) Lysis, mixing, and incubation: A biological sample is lysed, mixed with labeled antibodies, and incubated to allow couplex formation (targets bound by two different labeled antibodies). (3) Digital PCR amplification: During dPCR, thousands of partitions are simultaneously amplified, and specific fluorescent signals are generated. Couplex-containing partitions are detected as double-positive partitions (green). (4) Data analysis with PIQuant software: Raw dPCR data are uploaded and converted into molar concentrations of protein complexes using Poisson statistics.

**Supplemental Figure S2. Relationship between HTLV-1 proviral load and cytokine or viral gene expression.** Correlation analysis assessing the relationship between HTLV-1 proviral load (PVL) and (A–F) cytokine mRNA levels or (G) viral gene expression (Tax, HBZ) in CD8^+^-depleted PBMCs from HTLV-1 asymptomatic carriers (AC) and acute ATL patients. No significant correlation was observed between PVL and cytokine upregulation or viral gene expression, indicating that cytokine expression is independent of viral load in ATL.

**Supplemental Figure S3. Activation of pro-inflammatory cytokine promoters by Tax.** Luciferase reporter assays measuring activation of (A) IL-1β, (B) IL-2, (C) IL-4, (D) IL-17, and (E) CSF2 (GM-CSF) promoters by HTLV-1 Tax. HEK293T cells were co-transfected with luciferase reporter plasmids for the indicated cytokines and HTLV-1 Tax (HPX-Tax) or empty vector (Mock). 48 h after transfection, luciferase activity was measured. Tax expression was confirmed by immunoblotting, with nucleolin (C23) serving as a loading control. Data are representative of 4 independent experiments. Statistical analysis: one-way ANOVA with Dunnett’s multiple comparisons test (ns p > 0.05, *p < 0.01, **p < 0.001).

**Supplemental Figure S4. Chromatin immunoprecipitation analysis of Fra-2 occupancy at IL-2, TNF-α, and GM-CSF promoter regions.** ChIP-qPCR analysis of Fra-2 occupancy at NF-κB and AP-1 binding sites within the (A) IL-2, (B) TNF-α, and (C) GM-CSF (CSF2) promoters in ATL-2 cells. Fra-2 was immunoprecipitated using anti-Fra-2 antibody (G-5, sc-166102X), and DNA was analyzed by qPCR. Data are presented as fold enrichment relative to IgG control. Data are representative of 3 independent experiments. Statistical analysis: two-way ANOVA (*p < 0.01, **p < 0.001).

---

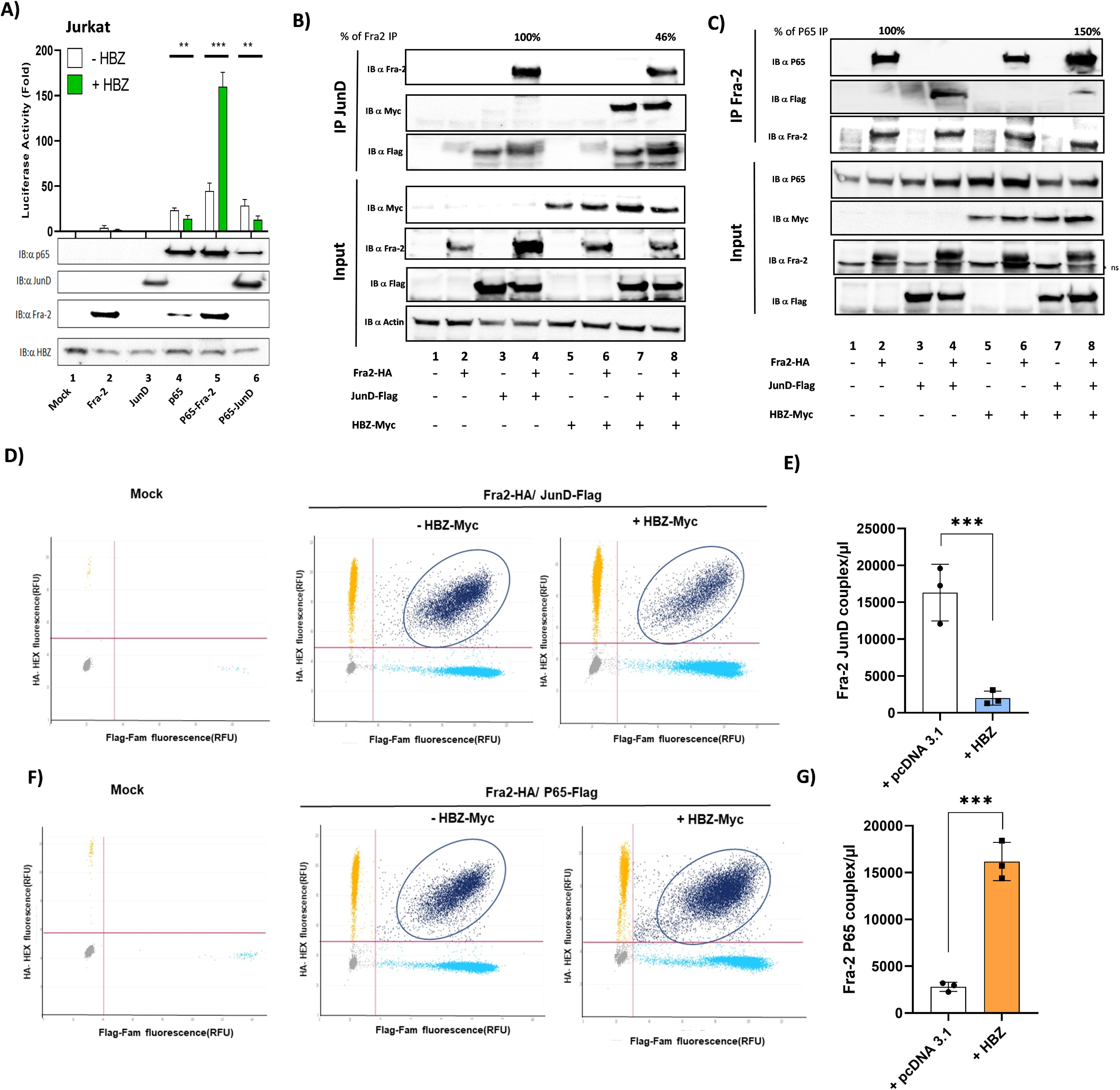

## Notes

### Competing Interest Statement

The authors have declared no competing interest.

### Funding

- Fondation pour la Recherche Médicale: DEQ20161136701
- Agence Nationale de la Recherche: 22-CE18-0019-01

