## supplementary table 1 for "Transcriptional rewiring of NF-κB by HBZ and Fra-2 drives inflammatory chemoresistance in Tax-negative adult T-cell leukemia/lymphoma"

**Supplemental Table S1 (Patient Demographics)**

| Group | n | Age (mean ± SD) | Sex (M/F) | HTLV-1 PVL (copies/10⁴ PBMCs) | ATL Subtype |
| --- | --- | --- | --- | --- | --- |
| Asymptomatic Carriers (AC) | 6 | 50 ± 12 | 3/3 | 10–50 | N/A |
| Acute ATL (Type I) | 6 | 60 ± 8 | 4/2 | 100–500 | Type I  Tax-negative |

**Supplemental Tables S2-RT-qPCR and dPCR primer sequences**

| Supplemental Table 1: Primers used for RT-qPCR | | | |
| --- | --- | --- | --- |
| Genes: |  | **Unique Assay ID:** | **Assay Design:** |
| IL1-α | Biorad | qHsaCID0016254 | Intron-spanning |
| IL1-β | Biorad | qHsaCID0022272 | Intron-spanning |
| IL-2 | Biorad | qHsaCID0015409 | Intron-spanning |
| IL-4 | Biorad | qHsaCID0013645 | Intron-spanning |
| IL-17 | Biorad | qHsaCID0015941 | Intron-spanning |
| TNF-α | Biorad | qHsaCED0037461 | Exonic |
| IFN-γ | Biorad | qHsaCED0043378 | Exonic |
| GM-CSF | Biorad | qHsaCED0048293 | Exonic |
| Genes: | **sequence** | |  |
| HPRT -FW | AAGGGCATATCCTACAACAAAC | |  |
| HPRT -Rev | GGTCAGGCAGTATAATCCAAAG | |  |
| Tax-FW | ACCAACACCATGGCCCA | |  |
| Tax-Rev | GAGTCGAGGGATAAGGAAC | |  |
| HBZ-Fw | ATGGCGGCCTCACCGTCGCAG | |  |
| HBZ-Rev | GGTCAGGCAGTATAATCCAAAG | |  |

**Supplemental Tables S3-ChIP primers,**

| Name | seq | Size amplicon (bp) |
| --- | --- | --- |
| NF-kB Binding sites |  |  |
| IL1A NF-kB FW | TAGAGATGGGGGCTTCACTATG |  |
| IL1A NF-kB REV | AACTAGGTGTGATGGCATGC | 121 |
| IL1B NF-kB FW | TTGGCCCTTCATTGTACCCA |  |
| IL1B NF-kB REV | CGTTGTGCAGTTGATGTCCAC | 89 |
| IL2 NF-kB FW | CCCCACCCCCTTAAAGAAAG |  |
| IL2 NF-kB REV | CCCCAAAGACTGACTGAATGG | 137 |
| TNF NF-kB1 FW | CAGAAGACCCCCCTCGGAATC |  |
| TNF NF-kB1 REV | GTTGGGGACACACAAGCATC | 77 |
| TNF NF-kB2 FW | GAGACAGAAGGTGCAGGGCC |  |
| TNF NF-kB2 REV | CAGCGGAAAACTTCCTTGGTG | 78 |
| CSF2 NF-kB FW | TCCCATGTGTGGCTGATAAGG |  |
| CSF2 NF-kB REV | CTTTCTCTCTGTGTACTGGGCTC | 144 |
| AP-1 Binding sites |  |  |
| IL1A AP-1A FW | CCCTGTTGAAGCTTTCAAGCC |  |
| IL1A AP1A Rev | GAGCTGCTATGGAGATGCAA | 154 |
| IL1A AP-1B FW | CATTTGCTAAGAGTCTGGTG |  |
| IL1A AP1B Rev | TGGAGATTGGGAGAAAGGAAG | 73 |
| IL1B AP-1 FW | TGGACATCAACTGCACAACG |  |
| IL1B AP-1 Rev | AGGGCAAGGAGTAGCAAAC | 95 |
| IL2 AP-1 FW | CCATTCAGTCAGTCTTTGGGG |  |
| IL-2 AP-1 REV | CTGGAAAAATATTATGGGGGTGTC | 148 |
| TNF AP-1A FW | CAAACACAGGCCTCAGGACTC |  |
| TNF AP-1A REV | AACTAGAACTGGGAGGGGCTTC | 97 |
| TNF AP-1B FW | GACCTGGTCCCCAAAAGAAATG |  |
| TNF AP-1B REV | GATTCCGAGGGGGGTCTTCTG | 117 |
| CSF2 AP1A FW | ACTCAGGAAGGTGGCTCCAGAG |  |
| CSF2 AP1A FW | ACTCAGGAAGGTGGCTCCAGAG | 79 |
| CSF2 AP1B FW | CTCACACTCAAGTCTCTCACAG |  |
| CSF2 AP1B REV | TCCCCATCCTGCCTGTAATC | 114 |
