## Supplementary figures and images for "Transcriptional rewiring of NF-κB by HBZ and Fra-2 drives inflammatory chemoresistance in Tax-negative adult T-cell leukemia/lymphoma"

### supplementary Figure 1

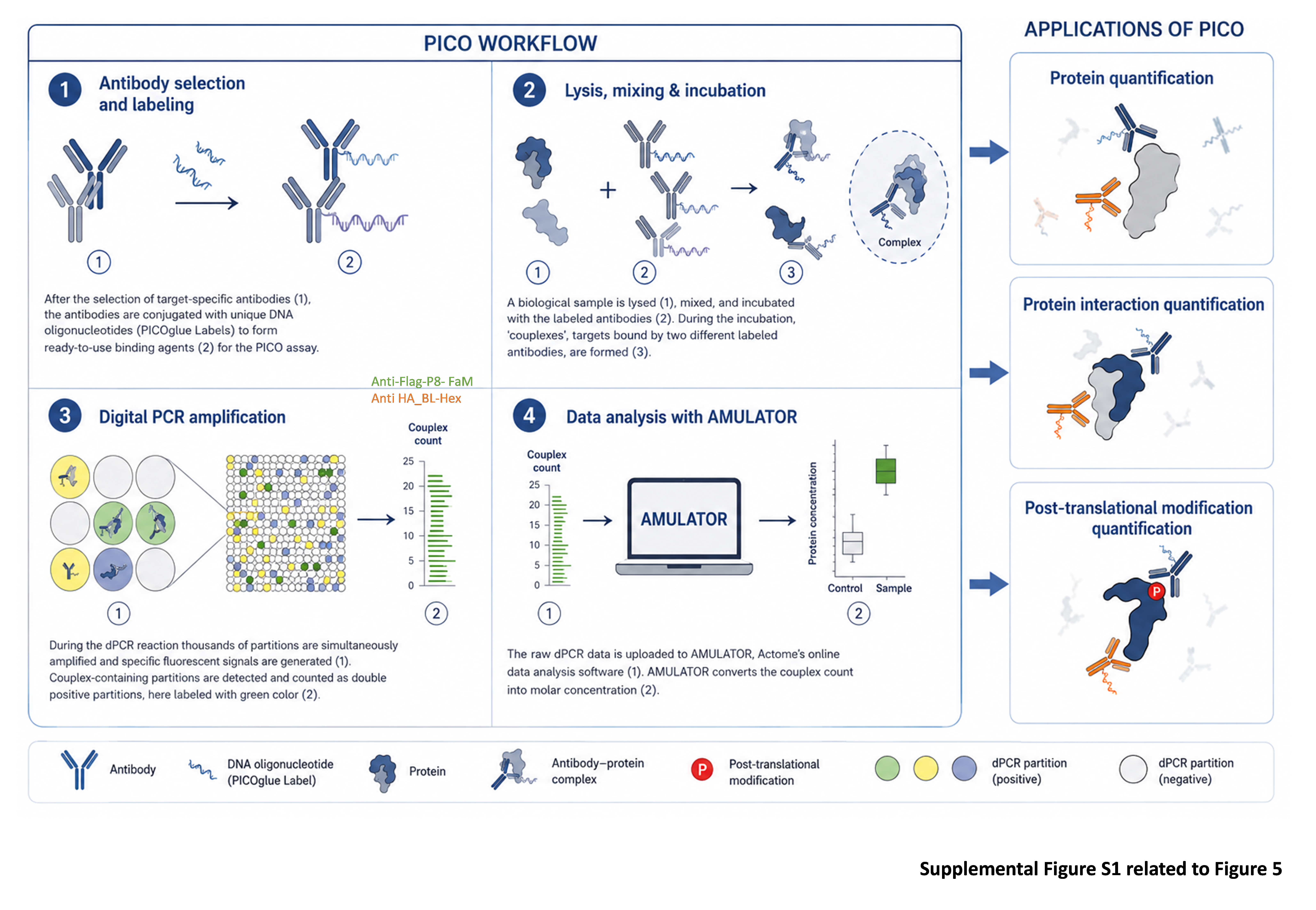

### supplementary Figure 2

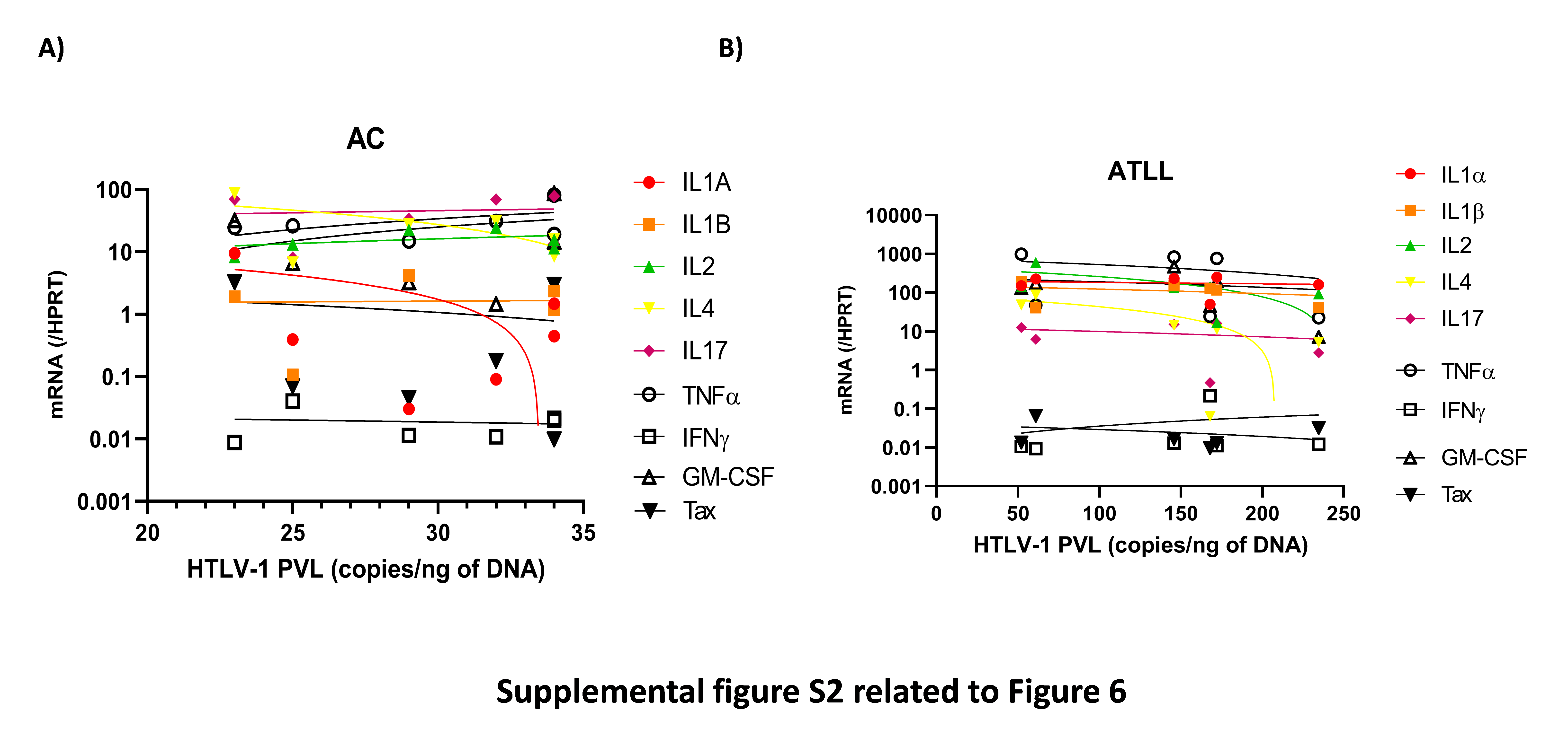

### supplementary Figure 3

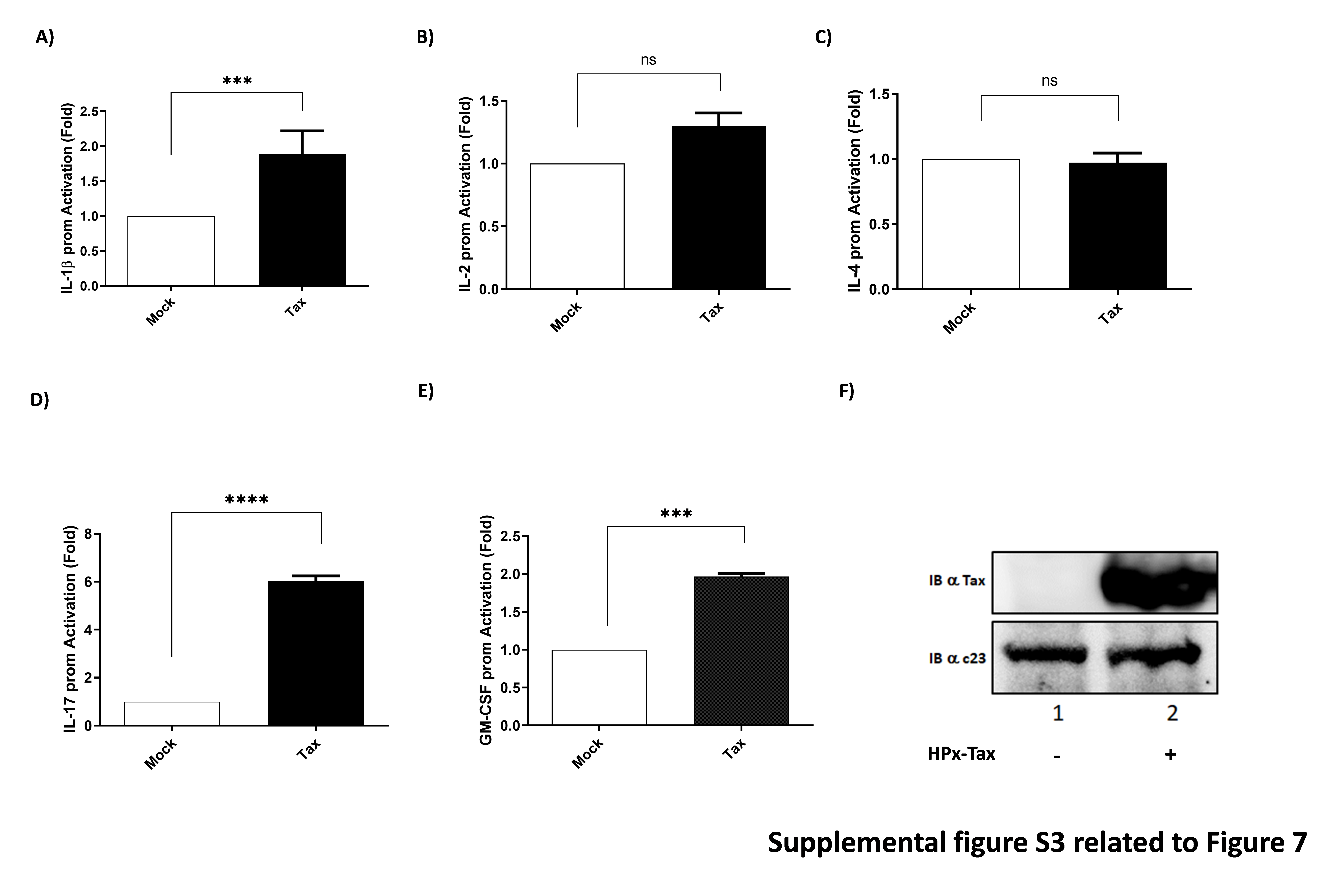

### supplementary Figure 4

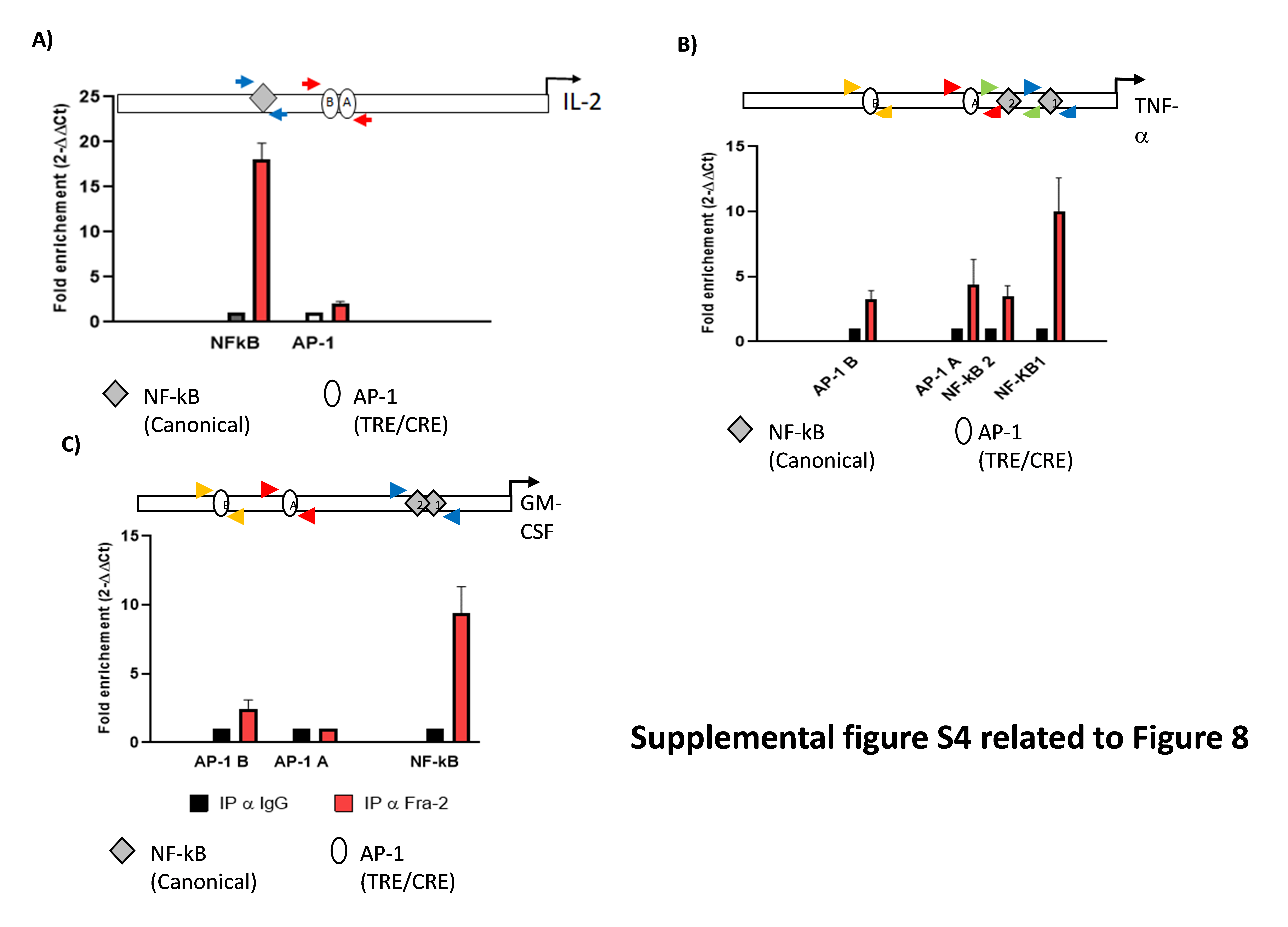
